# The C-terminal domain of RNase H and the C-terminus amino acid residue regulate virus release and autoprocessing of a defective HIV-1 possessing M50I and V151I changes in integrase

**DOI:** 10.1101/2022.10.23.513430

**Authors:** Tomozumi Imamichi, Qian Chen, Ming Hao, Weizhong Chang, Jun Yang

## Abstract

Previously, we reported that an HIV-1 variant containing Met-to-Ile change at codon 50 and Val-to-Ile mutation at codon 151 of integrase (IN), HIV(IN:M50I/V151I), was an impaired virus. Despite the mutations being in IN, the virus release was significantly suppressed (p < 0.0001) and the initiation of autoprocessing was inhibited; the mechanism of the defect remains unknown. In the current study, we attempted to identify the critical domains or amino acid (aa) residue(s) that promote defects in HIV(IN:M50I/V151I), using a series of variants, including truncated or aa-substituted RNase H (RH) or IN. The results demonstrated that virus release and the initiation of autoprocessing were regulated by the C-terminal domains (CTDs) of RH and IN. Further studies illustrated that Asp at codon 109 of RH CTD and Asp at the C terminus of IN induces the defect. This result indicated that the CTD of RH and IN in GagPol and particular aa positions in RH and IN regulated the virus release and the initiation of autoprocessing, and these sites could be potential targets for the developing new therapies.

## Introduction

Nascent viral particles of human immunodeficiency virus type-1 (HIV-1) released from HIV-1-producing cells are immature and uninfectious (1, 2). The particles contain Gag and GagPol polyproteins, regulatory viral proteins (Vif, Vpr, Vpu, Tat, Rev, and Nef), and two molecules of single-stranded genomic RNAs with host proteins (2–4). Gag and GagPol polyproteins and the regulatory viral proteins are encoded in a single genomic RNA. GagPol polyproteins are produced by a −1 nucleotide (nt) ribosomal frameshifting event during the translation of Gag protein (5). A ratio of translated Gag to GagPol is generally known as 20:1 in retrovirus (6). The maintenance of the ratio is important for HIV infectivity (7). Gag and GagPol translocate target the plasma membrane via the myristoylated Gly at the N terminus of the polyprotein, matrix protein (MA). Gag consists of MA, capsid (CA), nucleocapsid (NC), the spacer peptides p2 and p1, and p6 domains, and GagPol comprises MA, CA, NC, p6*, p2, p1, protease (PR), reverse transcriptase (RT), RNase H (RH) and integrase (IN) (1). Each domain is concatenated via sequences of PR cleavage site to form the polyprotein chains (1). Gag and GagPol assemble into a hexagonal lattice via its CA domain, recruiting viral proteins and the genomic viral RNAs. During assembly, the curved Gag lattices are formed corresponding to bending of the membrane at the site, followed by recruitment of the endosomal sorting complex III (ESCRT-III) components required for transport by the p6 domain of Gag and induces membrane scission and release of nascent virus particles (2, 8–11).

The nascent immature HIV-1 particles mature into infectious viruses through a sequential processing step of enzymatic cleavage of Gag and GagPol proteins by the mature PR (2, 3). The newly translated embedded PR in the GagPol polyprotein is not a fully functional enzyme. However, it is converted into a mature functional enzyme via self-cleavage in intramolecular manner (*cis*) (12); this step is called autoprocessing. Once the protein is converted to its active form, it cleaves the Gag and GagPol polyproteins at the cleavage sites and scissors each functional protein inside the virus particle during budding or after releasing (1). Autoprocessing is primed at the junction between p2 and NC as the initial cleavage of the same polyprotein chain with a strict regulation by the order of the cleavage sequence (12–14). The first digestion triggers a cascade of enzymatic reactions to release the functional mature PR. The autoprocessing step is suppressed by PR inhibitors (15, 16) or regulated by truncated IN (17–19), amino acid substituted IN (20, 21) or RH (21); thus, the autoprocessing step is considered a target for developing novel anti-HIV drugs.

A genome-wide association study of HIV genomic RNA demonstrated that a total of 14 non-synonymous single nucleotide polymorphisms (SNPs) were correlated with plasma viral load (VL) in anti-retrovirus treatment-naive HIV-infected patients (22). The mutations were located on CA, RH, IN, envelope, and Nef We previously investigated the roles of each SNP in viral fitness *in vitro* using a laboratory-adapted HIV clone, HIVNL4.3, as a parent strain and point mutagenesis was conducted to induce individual or a combination of the SNPs (21). We found that a mutant containing a Met-to-Ile change at codon 50 of IN (IN:M50I) was impaired. The mutant suppressed the virus release to 0.3% of wild-type HIV_NL4.3_ (WT), lacking GagPol autoprocessing (21, 23), indicating that IN regulates autoprocessing, consistent with previous reports (17–20). The defect was rescued by the co-existing other VL-associated SNPs, Ser-to-Asn change at codon 17 of IN (IN:S17N) (21). Interestingly, an Asn-to-Ser change at codon 79 of RH (RH:N79S) also restored virus release, autoprocessing and replication competence of the defective M50I virus (21). These results indicated that RH is also involved in virus release and autoprocessing, thereby affecting virus fitness.

We investigated a population of HIV-containing the IN:M50I mutation in patients participating in NIAID clinical trials. Surprisingly, the plasma VL in patients infected with HIV containing IN:M50I without the compensatory mutations (either IN:S17N or RH:N79S) was 1207–201,353 copies/ mL (23), suggesting that the clinical isolates containing IN:M50I were not impaired in the virus released from HIV-producing cells, unlike our previous *in vitro* study. To clarify this discrepancy between the *in vitro* and the *in vivo* data, we performed a comparative analysis of the RNA genome of the entire *pol* region. We then found that the recombinant HIV(IN:M50I) virus contained a Val-to-Ile mutation at codon 151 of IN (IN:V151I) in the backbone (23). IN:V151I is an endogenous aa substitution in HIV_NL4.3_ (24), and is known as a polymorphic mutation associated with drug resistance (25–29). Given the reports that a recombinant consensus B HIV_NL4.3_ IN protein-containing M50I and V151I is functional (30), while our recombinant virus possessing the mutations was defective in virus release and autoprocessing, we hypothesized that backbone mutations in the HIV(M50I/V151I) may regulate virus release, autoprocessing and replication *in vitro* study.

In the current study, we further investigated the mechanism of the defect in virus release and autoprocessing in HIV(M50I/V151I) and report that the C-terminal domains of RH and IN are associated with the deficiency. These results provide a potential target for the development of anti-HIV drugs/therapy.

## 2. Materials and Methods

### 2.1. Ethics Statement

Approval for these studies, including all sample materials and protocols, was granted by the National Institute of Allergy and Infectious Diseases (NIAID) Institutional Review Board, and participants have informed a written consent prior to blood being drawn. All experimental procedures in these studies were approved by the National Cancer Institute at Frederick and Frederick National Laboratory for Cancer Research (the protocol code number: 16–19, approval data: 6 January 2017).

### 2.2. Cells

Peripheral blood mononuclear cells (PBMCs) were isolated from healthy donors’ apheresis packs (NIH blood bank) using a lymphocyte separation medium (ICN Biomedical, Aurora, OH, USA) (21, 23), CD4(+) T cells were purified from PBMCs using CD4 MicroBeads (Miltenyi Biotec, Auburn, CA, USA) according to the manufacturer’s instructions. The purity of the cell types was at least 90%, based on flow cytometric analysis. Cell viability was determined using the trypan blue (Thermo Fisher, Waltham, MA, USA) exclusion method. HEK293T cells were obtained from ATCC (Manassas, VA, USA) and maintained in complete D-MEM (Thermo Fisher) supplemented with 10 mM 4-(2-hydroxyethyl)-1-piperazineethanesulfonic acid (HEPES) pH 7.4 (Quality Biological, Gaithersburg, MD, USA), 10% (v/v) fetal bovine serum (FBS; Thermo Fisher), and 50 μg/mL gentamicin (Thermo Fisher) as previously described (31).

### 2.3. Construction of Plasmids Encoding HIV Variants

Plasmids encoding HIV variants were constructed by inducing the point mutagenesis on pNL4.3 (32) (the plasmid was obtained from M. Martin through the AIDS Research and Reference Reagent Program, National Institute of Allergy and Infectious Diseases, National Institutes of Health). A point mutagenesis was performed using the QuickChange Lightning kit (Agilent

Technologies, Santa Clara, CA, USA) with mutagenesis primers (Table S1) as previously described (21). RH or IN-deleted constructs were created by invert PCR using Pfu II Taq polymerase (Agilent Technology) with deletion primers (Table S1). Mutagenesis was confirmed by Sanger DNA sequencing using the BigDye terminator v.3 (Thermo Fisher) with SeqStudio Genetic Analyzer (Thermo Fisher), and plasmids were maintained in Stbl3 E. coli (Thermo Fisher). The plasmid purification was carried out using the EndoFree Plasmid Maxi Kit (Qiagen, Germantown, MD, USA).

### 2.4. Recombinant HIV-1 Viruses

Recombinant HIV-1 variants were prepared by transfection of the pNL4.3 mutants into HEK293T cells using TransIT-293 (Mirus, Houston, TX, USA) and Opti-MEM I medium (Thermo Fisher) following a method previously reported (21). Culture supernatants were collected at 48 h after transfection and then filtrated through 0.45 μm pore size filter membranes (MiliporeSigma, Burlington, MA, USA). Virus particles in the filtrate were pelleted by an ultra-centrifugation on 20% (w/v) sucrose (MiliporeSigma) in 10 mM HEPES-150 mM NaCl buffer (pH 7.4) as previously described (21, 33). The pelleted virus particles were washed with PBS and then resuspended in PBS in 1/100 vol of supernatants. The virus stocks were stored at −80 °C until use. Concentration of HIV p24 antigen in each stock was determined by a p24 antigen capture kit (PerkinElmer, Waltham, MA, USA) and the concentration of total virus proteins were determined by a BCA protein assay kit (Thermo Fisher) (31, 33).

### 2.5. HIV Replication Assay

Replication capability of each variant were determined using primary CD4(+) T cells as previously described (21). CD4(+) T cells were stimulated with 5 μg/mL phytohemagglutinin (PHA; MiliporeSigma) in complete RPMI-1640 (Thermo Fisher) supplemented with 10 mM HEPES, 10% (v/v) FBS, and 50 μg/mL gentamicin (RP10). The PHA-stimulated CD4(+) T cells (10 × 10^6 cells) were infected with 10 ng of p24 of each HIV variant at 10 × 10^6 cells/mL in RP10 for two hours at 37 °C. The infected cells were washed with RP10, and then cultured at 1 × 10^6 cells/mL in the medium in the presence of 20 units/mL of recombinant IL-2 (MiliporeSigma) for 14 days at 37 °C in T25 flasks (21, 23). Half of the cell-free culture supernatants were exchanged with fresh RP10 with 20 units/ml of IL-2 every 3 or 4 days of incubation. HIV-1 replication activity was determined by measuring p24 antigen levels in the culture supernatants using the p24 antigen capture assay (PerkinElmer, Boston, MA, USA) (23).

### 2.6. Western Blotting

To confirm autoprocessing of Gag and GagPol polyproteins in HIV virions, Western Blotting (WB) was performed as previously described (21, 23). Due to abnormal sizes of M50I mutant (the diameters of the mutant particles were 190~300 nm, while those of Wt were 110~130 nm, and in this current study, the mutant previously used is renamed mNL(IN:M50I) as described in Section 3.1 below) with a defective autoprocessing of Gag and GagPol polyproteins (lacking of the cleaved mature forms of MA (p17), CA (p24), PR, RT, and IN in the virus particles) (21), we could not find an appropriate internal control for WB. Thus, we used 1 μg of total viral protein to demonstrate an impaired autoprocessing as previously described (21, 23). WB was performed using Rabbit polyclonal anti-Anti-HIV1 p55/p24/p17 antibody (Cat# ab63917, Abcam, Waltham, MA, USA) and anti-protease antibody (Cat# ab211627 Abcam, Cat# SKU: 65-018, As One International, Santa Clara, CA, USA), Protein bands were detected by using the ECL Prime Western Blotting Detection Reagent (MiliporeSigma) with the Azure 300 (Azure Biosystem (Dublin, CA, USA) (21).

### 2.7. Structure analysis

The HIV Cryo-EM Structure of the PR-RT structure (PDB accession #: 7sjx) (34) was downloaded from the PDB database (https://www.rcsb.org) into PyMOL (https://pymol.org/). The chain A in the structure was colored with limon (RH part was colored cyan) and the chain B was colored with olive. The RH:N79N and RH:D109 is only visible in chain A (Chain A:D630) colored as red and blue, respectively. The distance of alpha carbon between the RH:N79 and RH:D109 and was calculated with the command: distance chain A and i. 600 and n. CA, chain A and i. 630 and n. CA.

### 2.8. Statistical Analysis

Intergroup comparisons were performed by one-way analysis of variance (ANOVA) with multiple comparison analysis or Student’s t-test using GraphPad Prism 9 (GraphPad, San Diego, CA, USA). P values less than 0.05 were considered statistically significance (* P < 0.05, ** P < 0.01, *** P < 0.001, **** P<0.0001, P > 0.05 was considered not significant (ns).

## 3. Results

### 3.1. The C-terminal domain of RNase H in the GagPol polyprotein regulates the virus release and autoprocessing defects in HIV(IN:M50I/V151I)

We reported that RH:N79S or IN:S17N restored the inhibition of virus release and autoprocessing of HIV(IN:M50I/V151I) (21). In the current study, we attempted to determine whether the domains containing the changes directly regulate the virus release and the initiation of the autoprocessing. RH consists of four α-helixes (α1 ~ α4 helixes) and five β-sheets (β1~ β5 sheets), and the codon 79 is located in the middle of the α3 helix (Figure 1A, Supplemental Figure S1) and forms the active pocket with a highly conserved DEDD motif (the motif is composed of codon D3 on the β1-sheet, codon E38 on the α1-helix, codon D58 on the loop between the β3-sheet and the α2-helix, D109 on the α4-helix) which coordinates two divalent cations (Mn^2+^ or Mg^2+^) (35) required for hydrolyzing the HIV genomic RNA substrate. Although the compensatory mutation, RH:N79S is located on the α3 helix, which is outside of the metal binding pocket with 15.3 ~28.3Å distance from the activation site (Supplemental Figure S1 and S2), it rescued virus release and initiated autoprocessing followed by virus replication (21). Thus, we thought RH:N79S may trigger a structure change, subsequently induce recovery; other domains/regions of RH may directly associate with the recovery. We decided to determine how the domain(s) influence the virus maturation and constructed a series of HIV(M50I/V151I) variants containing different lengths of RH (Figure 1B). To evaluate the processing function of PR, we maintained the PR cleavage sequence at the junction between RH and IN in all variants; all mutants retained five amino acid (aa) sequences of the PR cleavage site at the C terminal of RH. Each plasmid construct was transfected into HEK293T cells. The released virus particles in the culture supernatants were collected by centrifugation and then resuspended in PBS or fresh culture medium, as described in the Materials and Methods. The amounts of released virus particles were quantified using a p24 antigen capture assay. The processing of Gag and GagPol in the particles was analyzed by western blot (WB) assays using anti-Gag or anti-PR antibodies. Consistent with our previous results, the amounts of HIV(IN:M50I/V151I) released were suppressed to 0.23 ± 0.02% (n=5) (p<0.0001) compared to HIV WT (Figure 1C) and restored to 64 ± 0.1 (n=4) % (p>0.05) of HIV(WT) in the presence of RH:N79S. The GagPol processing was also rescued (Figure 1D and 1E). All variants lacking the α4 helix domain could recover the virus release to 40~60% of WT (Figure 1C, Supplemental Table S2). The processing was recovered, and the mature form of PR was detected in all mutants, even in the absence of the RH:N79S (Figure 1D and 1E). The result indicated that the α4 helix domain, rather than the α3 helix domain, played a key role in regulating the virus release and the autoprocessing in HIV(IN:M5I/V151) and it negatively regulates them.

**Figure 1.**
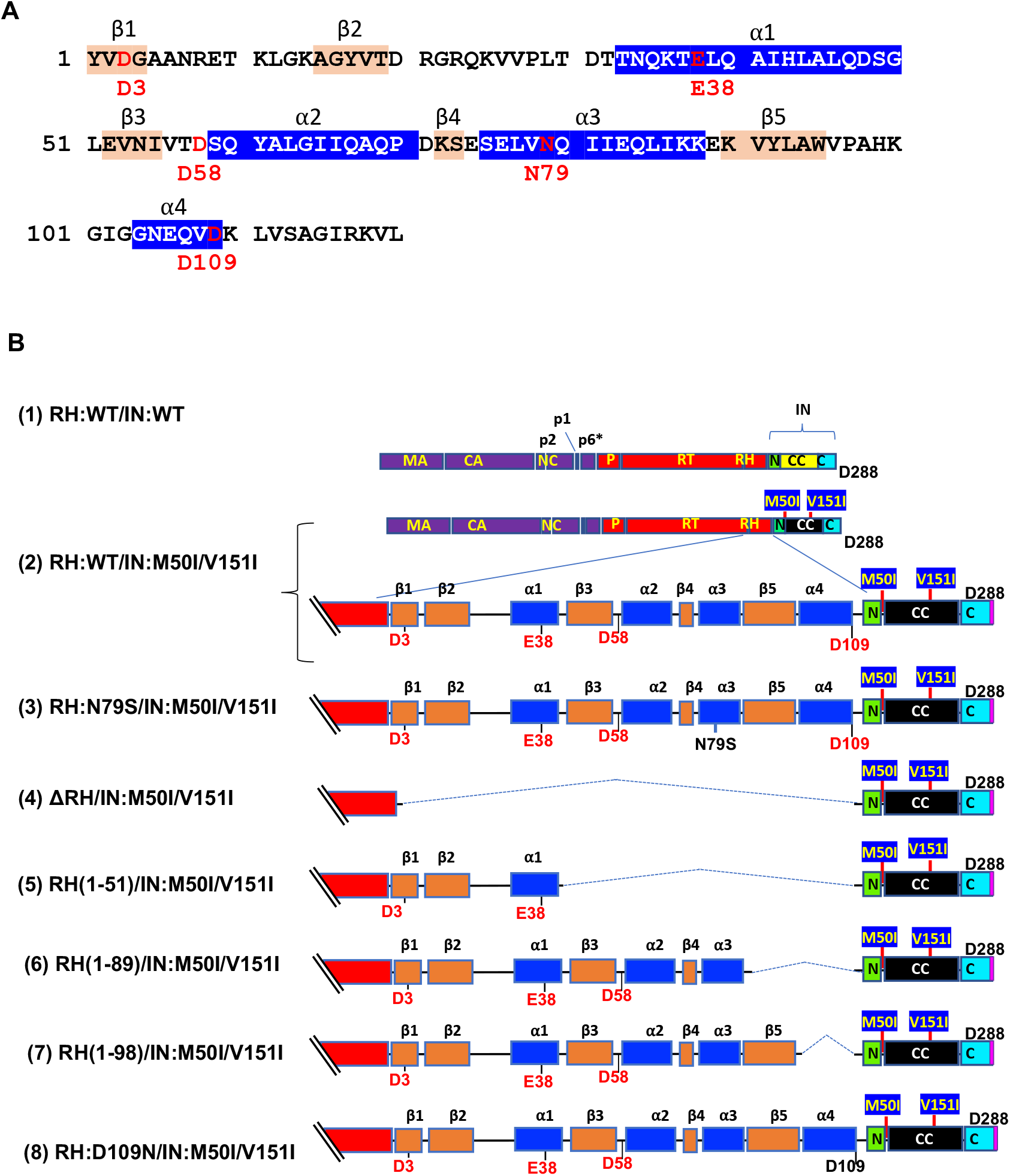
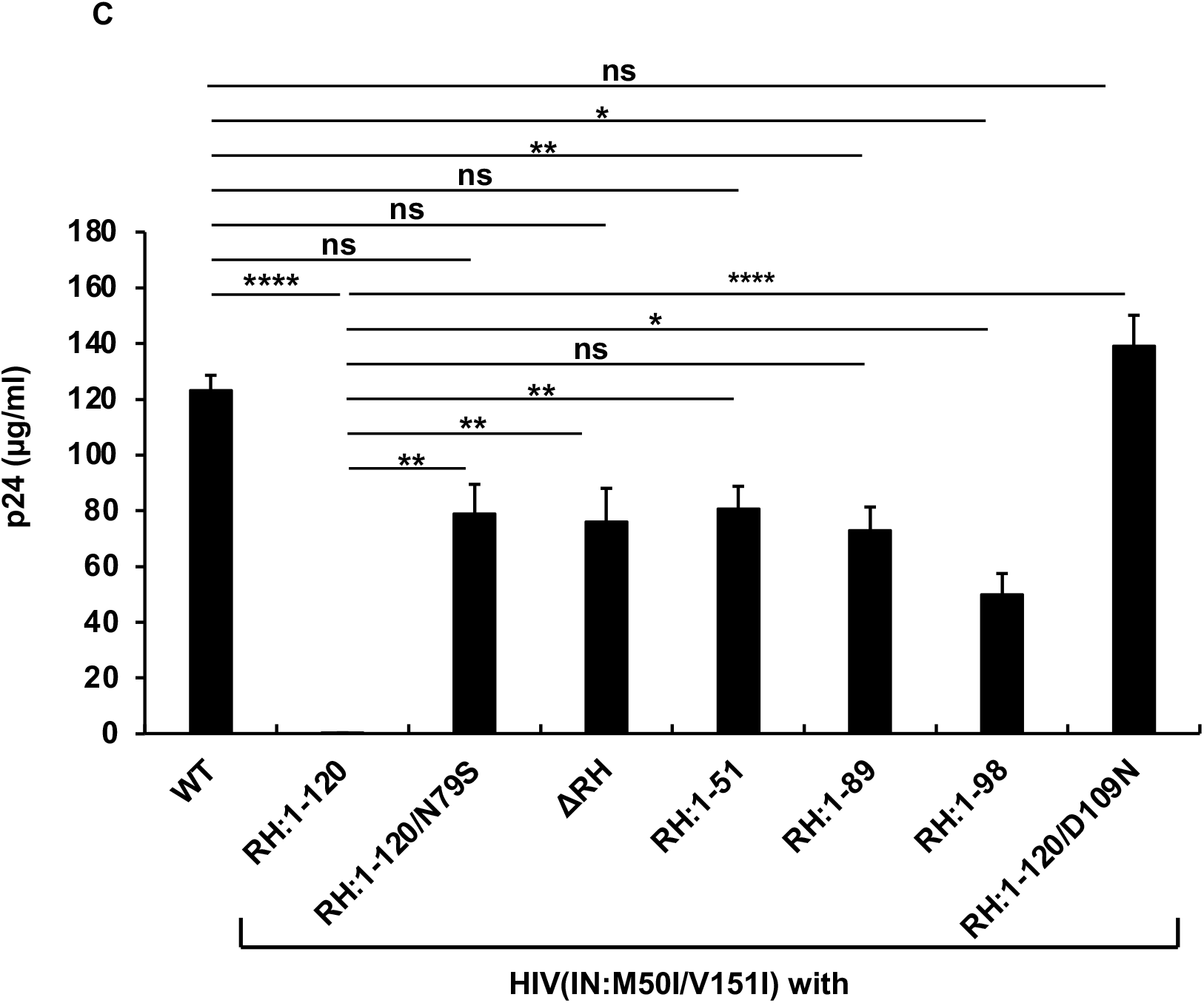
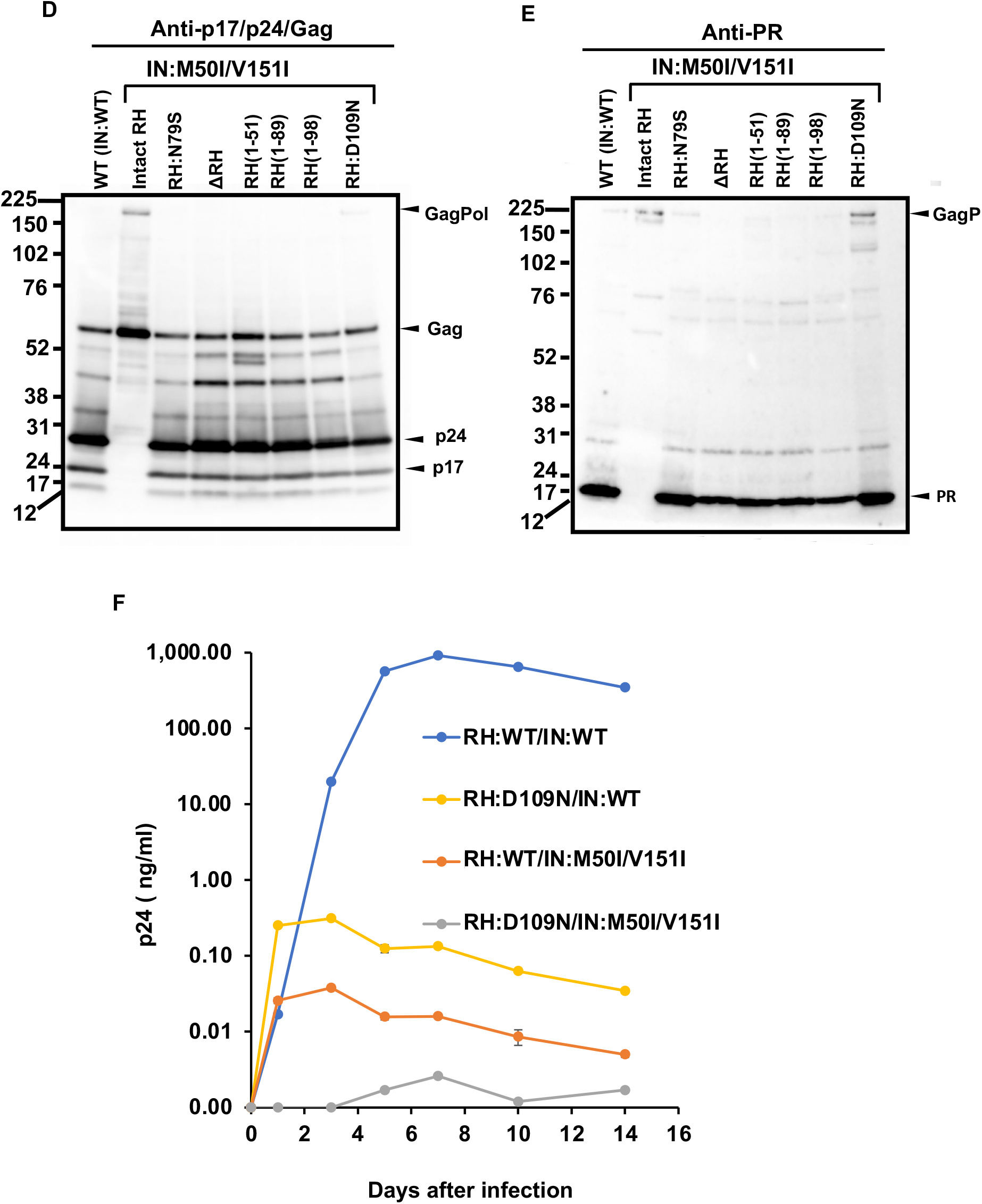
The C-terminal domain of RH regulates virus release and autoprocessing. (A) Amino acid sequence of RH. Amino acid sequence of RH of HIVNL4.3 (32) was obtained from GenBank (accession #: AF324493). The alpha helices are colored in blue, and the beta strands are colored in wheat; codons associated with the active sites (D3, E38, D58, and D109) and mutation position 79 (N79) are labeled and colored in red with numbering. (B) Diagrams of HIV mutants containing mutations in RH. The diagram indicates the structure of GagPol(WT) and variants. The numbers above the diagrams indicate codon positions in RH. Each mutant was created by inverse PCR or point mutagenesis as described in the Materials and Methods. MA: matrix protein, CA: capsid protein, NC: nucleocapsid protein, P: protease, RT: reverse transcriptase, RH: RNase H, IN: integrase, N: the N terminal domain of IN CC: the catalytic core domain of IN, C: the C terminal domain of IN, α1 ~ α4 and β1 ~ β5 are corresponding to domains of RH in Figure 1A. (C) ach construct was transfected into HEK293T cells. Transfection supernatants from HEK293T cells were collected, and then viral particles were pelted as described in the Materials and Methods. Each viral pellet was resuspended in 1/100 of the transfection supernatants in complete culture media or PBS. p24 concentration in the suspension was determined by measuring HIV p24 antigen by ELISA. Data indicates mean+ SE (n=5), and statistical analysis was conducted using one-way ANOVA (Prism). (D and E) Virus particles are isolated using ultracentrifugation as described in Materials and Methods, and 1 μg of viral lysates was subjected to WB. Gag/GagPol cleaved products and PR were detected by rabbit polyclonal anti-p17/p24/gag (D) and anti-PR (E), as described in the Materials and Methods. (F) PHA-stimulated primary CD4(+) T cells from healthy donors were infected with 10 ng p24 amounts of HIV(WT) or variants as described in Materials and Methods. The infected cells were cultured for 14 days with changing medium every 3 to 4 days. HIV replication was monitored using a p24 antigen capture kit. Representative data from three independent assays are presented as means ± standard deviations (SD) (n = 3).

The α4 helix domain is composed of six aa (Gly-Asn-Glu-Gln-Val-Asp, Figure 1A) containing RH:D109. The residue forms a metal-binding pocket in RH with other three residues (D3, E38, D58) (35). To define key amino acid residues in the domain, we performed a population analysis using HIV sequence data from LANL, Stanford drug resistance database, and NIAID HIV sequence data. The population analysis indicated that RH:D109 is highly conserved among the databases (LANL database: 99.62%, Stanford database: 100%, NIAID: 100%, Supplemental Table S3) (35); thus, we became interested in defining how RH:D109 affects the defective virus. The mutation Asp-to-Asn change at codon 109 (RH:D109N) was induced in HIV(IN:M50I/V151I), and then the amount of released virus and Gag/GagPol processing were analyzed. Interestingly, the mutation comparably restored virus release (Figure 1C, Supplemental Table S2) and the autoprocessing with HIV(WT) (Figure 1D and 1E); however, virus replication was defective (Figure 1F, Supplemental Figure S3), due to lack the functional metal binding pocket.

The defect of HIV(IN:M50I/V151I) in virus release was partially rescued in the absence of RH (Figure 1C), implicating that the RH domain in the GagPol negatively regulates the virus release in the defective virus. To define whether the RH domain affects virus release of HIV(IN:WT), the RH domain-deleted HIV(IN:WT), HIV(ΔRH/IN:WT) was created, and the virus release was assessed. If the RH domain negatively regulates the virus release, we expected that the release of HIV(ΔRH/IN:WT) might be increased; however, the mutant demonstrated 53± 11% (n= 3, p<0.05) of HIV(WT) (Figure 2A), indicating that the RH domain positively regulates virus release of HIV(WT). WB demonstrated that HIV(ΔRH/IN:WT) was able to process Gag and GagPol (Figure 2B and 2C), thus, RH domain is involved in virus release.

**Figure 2.**
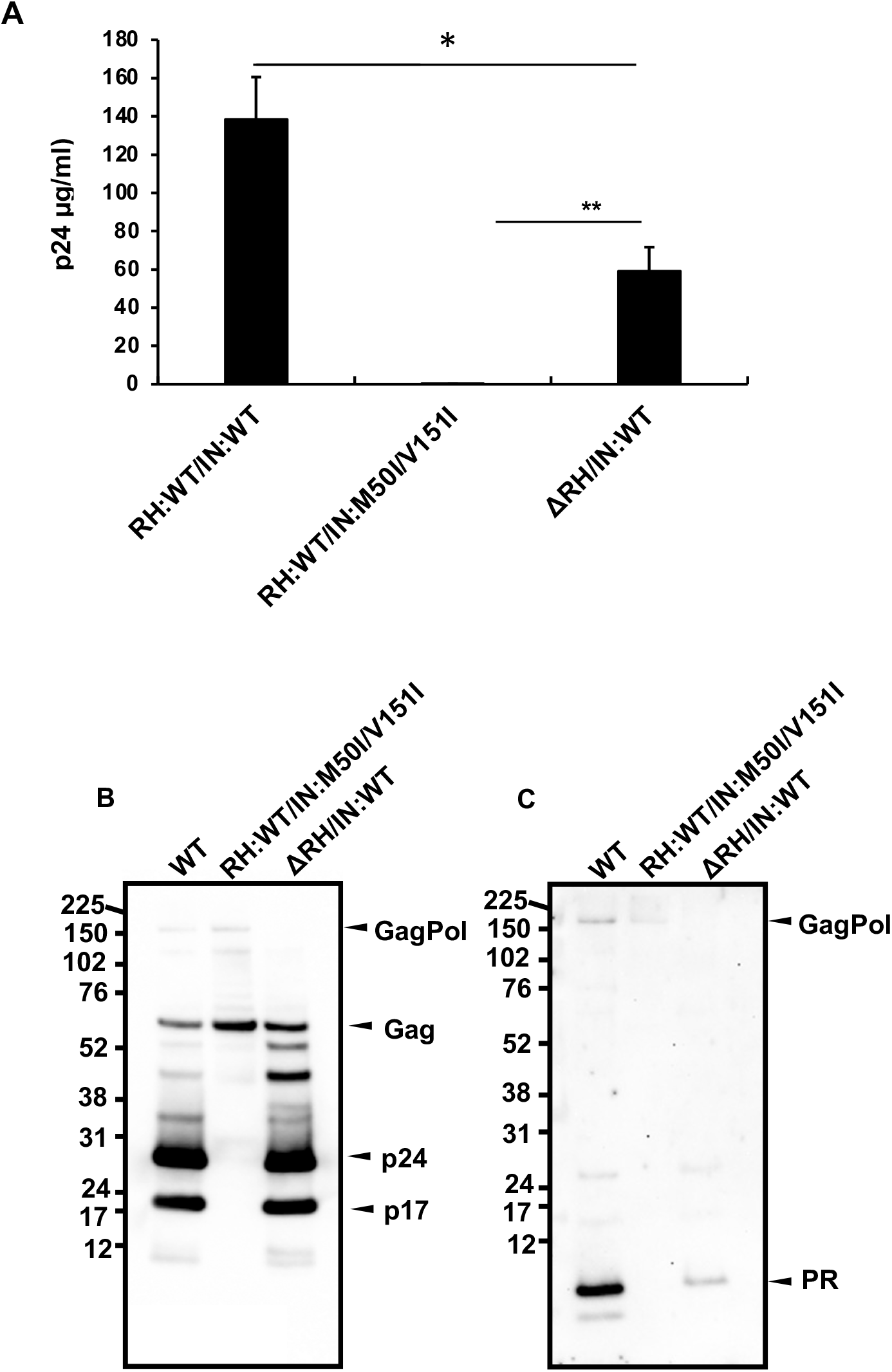
The C-terminal domain of RH regulates virus release and autoprocessing in HIV(IN:WT) (A, B, and C) Each construct was transfected into HEK293T cells. Transfection supernatants were collected, and viral particles were pelted as described in the Materials and Methods. Each viral pellet was resuspended in 1/100 of the transfection supernatants in complete culture media or PBS. (A) P24 concentration in the suspension was determined by measuring HIV p24 antigen by ELISA. Data indicate means ± SE (n=5) statistical analysis was conducted using one-way ANOVA (Prism). (B and C) Virus particles are isolated as described in Materials and Methods, 1 μg of viral lysates was subjected to WB. Gag/GagPol cleaved products and PR were detected by rabbit polyclonal anti-p17/p24/Gag (B) and anti-PR antibody (C),

### 3.2 Amino acid residue at the C-terminal of integrase (IN) Asp at 288 is a key player in the M50I/V151I defect in autoprocessing and virus release

As shown above, RH:D109N rescued the defects in autoprocessing and virus release in the context of HIV(IN:M50I/V151I). This implicated that the residue or the α4 helix domain of RH negatively regulates both the virus release and the initiation of the autoprocessing in the context. To further understand the suppression mechanism in the presence of RH:D109 and the role of domains of IN in inducing the defect, we constructed a series of variants (Figure 3A). IN is composed of three domains, the N-terminal domain (NTD), the catalytic core domain (CCD), and the C-terminal domain (CTD) (36–38). Codon 50 is located at the linker/hinge region between NTD and CCD, which is 21~25 angstrom away from the activation site of IN (Supplemental Figure S4). In contrast, codon 151 exists in CCD and is a neighbor of acidic residues in the active pocket. Thus, we focused on CTD to construct the mutants to maintain the M50I and V151I effect. Then virus release and autoprocessing were analyzed.

**Figure 3.**
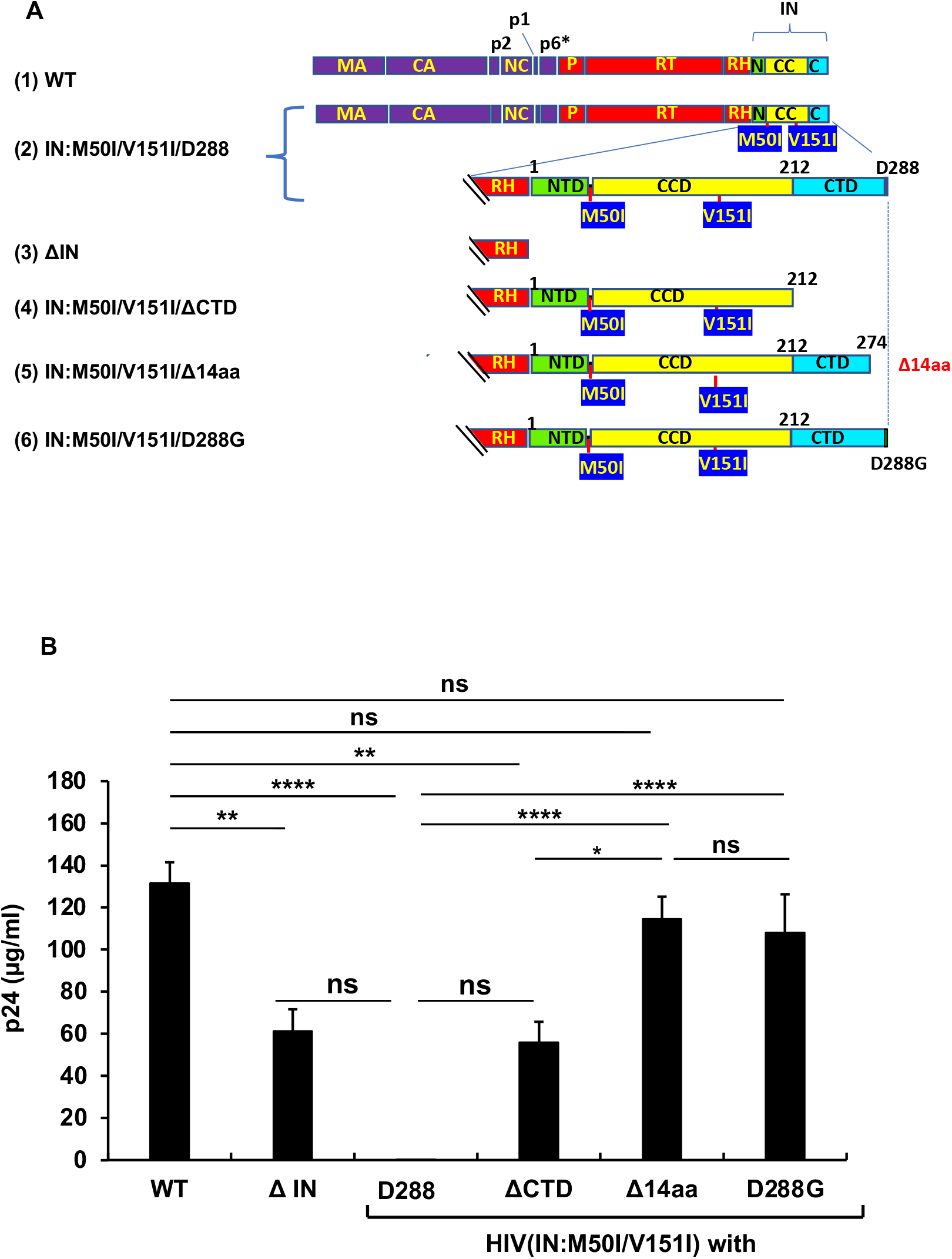
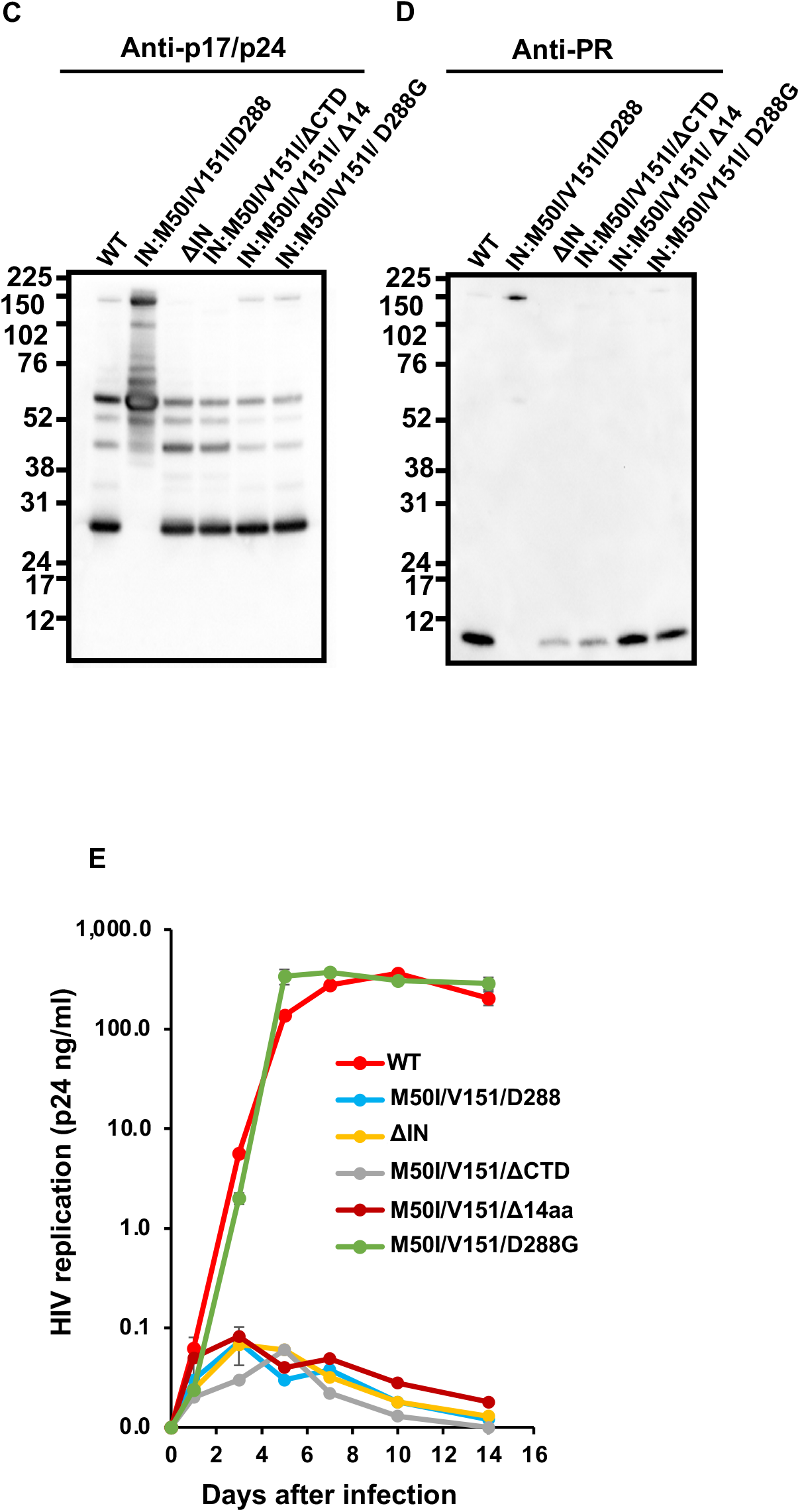
The C-terminal domain of IN regulates virus release and autoprocessing. (A) Diagrams indicate the structure of GagPol(WT) and variants. The numbers above the diagrams indicate codon position in IN. Each mutant was created by inverse PCR or point mutagenesis as described in Materials and Methods. MA: matrix protein, CA: capsid protein, NC: nucleocapsid protein, P: protease, RT: reverse transcriptase, RH: RNase H, IN: integrase, N: the N terminal domain of IN CC: the catalytic core domain of IN, C: the C terminal domain of IN. (B) Each construct was transfected into HEK293T cells. Transfection supernatants from HEK293T cells were collected, and viral particles were pelted as described in the Materials and Methods. Each viral pellet was resuspended in 1/100 of the transfection supernatants in complete culture media. Viral concentration was determined by measuring HIV p24 antigen by ELISA. Data indicate means ± SE (n=5), and statistical analysis was conducted using one-way ANOVA (Prism). (C and D) Virus particles are isolated as described in Materials and Methods, and 1 mg of viral lysates was subjected to WB. Gag and GagPol cleaved products and PR were detected by anti-p17/p24/gag (C) and anti-PR (D). (E) PHA-stimulated primary CD4(+)T cells from healthy donors were infected with 10 ng p24 amounts of HIV(WT) or variants containing mutations as described in Materials and Methods. The infected cells were cultured for 14 days with the medium change every 3 to 4 days. HIV replication was monitored using a p24 antigen capture kit. Representative data from two independent assays are presented as means ± standard deviations (SD) (n = 3).

The virus release of the mutant lacking the entire IN (ΔIN) was suppressed to 49.7± 8.5 % of that of WT (p<0.05, n=4) (Figure 3B), and WB results demonstrated that it partially suppressed autoprocessing (Figure 3C and 3D). Therefore, consistent with previous reports (7, 9), IN is involved in regulating autoprocessing. A mutant lacking the entire CTD, (HIV(M50I/V151I/ΔCTD) partially restored the virus replication (45.5± 7.9 % of WT, p<0.05, n=4) and autoprocessing (Figure 3C and 3D). Of interest, a mutant without the C-terminal tail of 14 aa of IN, HIV(IN:M50I/V151/Δ14aa) restored the virus release and processing to comparable levels of WT (Figure 3B, 3C and 3D) comparable to WT, indicating that the last 14 aa sequence (Met-Ala-Gly-Asp-Asp-Cys-Val-Ala-Gly-Arg-Gln-Asp-Glu-Asp) regulates both autoprocessing and virus release of HIV(IN:M50I/V151I) in the presence of RH:D109.

We also assessed the capability of HIV replication and compared its activity among mutants. It is reported that variants lacking the last 15~17 aa residues at the tail domain diminished enzymatic function in IN and infectivity (39, 40); consistent with the reports, all mutants lacking IN domains did not indicate a detectable level of HIV replication (Figure 3E).

We considered how the tail restored the autoprocessing even in the presence of RH:D109. Crystal structural analysis of the full length of IN has not yet been successful because of the flexibility, instability of the tail and its poor solubility under low salt conditions (34, 36, 41–43). It was assumed that the tail of IN:M50I/V151I might directly (e.g., structure hindrance) or indirectly (e.g., interaction with host proteins) influence the initiation of the autoprocessing at the junction of p2 and NC. If an unknown factor was involved in the defect, we thought the last amino acid residue might critically function rather than the middle of amino acid residues in the tail. Asp is an acidic amino acid; thus, we thought the charge or size of residue might influence the defect. To address the hypothesis, we chose an aa change at 288 from Asp to Gly, the smallest aa side residue without polar, as a pilot experiment. Using site mutagenesis, we constructed a mutant, HIV(IN:M50I/V51I/D288G), and then, virus release, autoprocessing, and virus replication assay were carried out. Surprisingly, the variant demonstrated indistinguishable activities in the virus release (Figure 3B, Supplemental Table S4), autoprocessing (Figure 3C and 3D), and virus replication (Figure 3E, Supplemental Figure S5), compared to WT. These data indicated that the last amino acid of the tail of IN regulates viral infectability in the IN:M50I/V151I mutant setting with RH:D109.

The D288G change alters the net charge and size of the side-residue at the tail. The LANL, Stanford, and NIAID HIV sequence databases demonstrated Asn at 288 in IN (IN:D288) is a relatively conserved in HIV sequences (the percentages of population of Asp at 288 in LANL, Stanford, and NIAID databases are 93.7%, 97.85 %, and 96.12%, respectively) (Supplemental Table S5); thus, to further define a correlation between the side residue and virus release/autoprocessing/virus replication, we produced a series of HIV(IN:M50I/V151I) mutants containing amino acid changes at codon 288. Neutral, basic, and hydrophobic amino acid changes were chosen from mutations detected in clinical isolates (Supplemental Table S5). In addition, to address the role of a polar side chain at 288, D288Y was also induced. The virus release of mutants containing D288K, D228A, D228E, or D288Y was restored to a comparable level of HIV(WT). A mutant containing D288N restored the virus release to only 50% of HIV(WT) (p<0.05) (Figure 4A); however, the processing levels in all mutants were similar to that of WT (Figure 4B and 4C) and virus replication was rescued in all mutants (Figure 4D), indicating that the last amino acid Asp at 288 may play a key role in regulating virus release.

**Figure 4.**
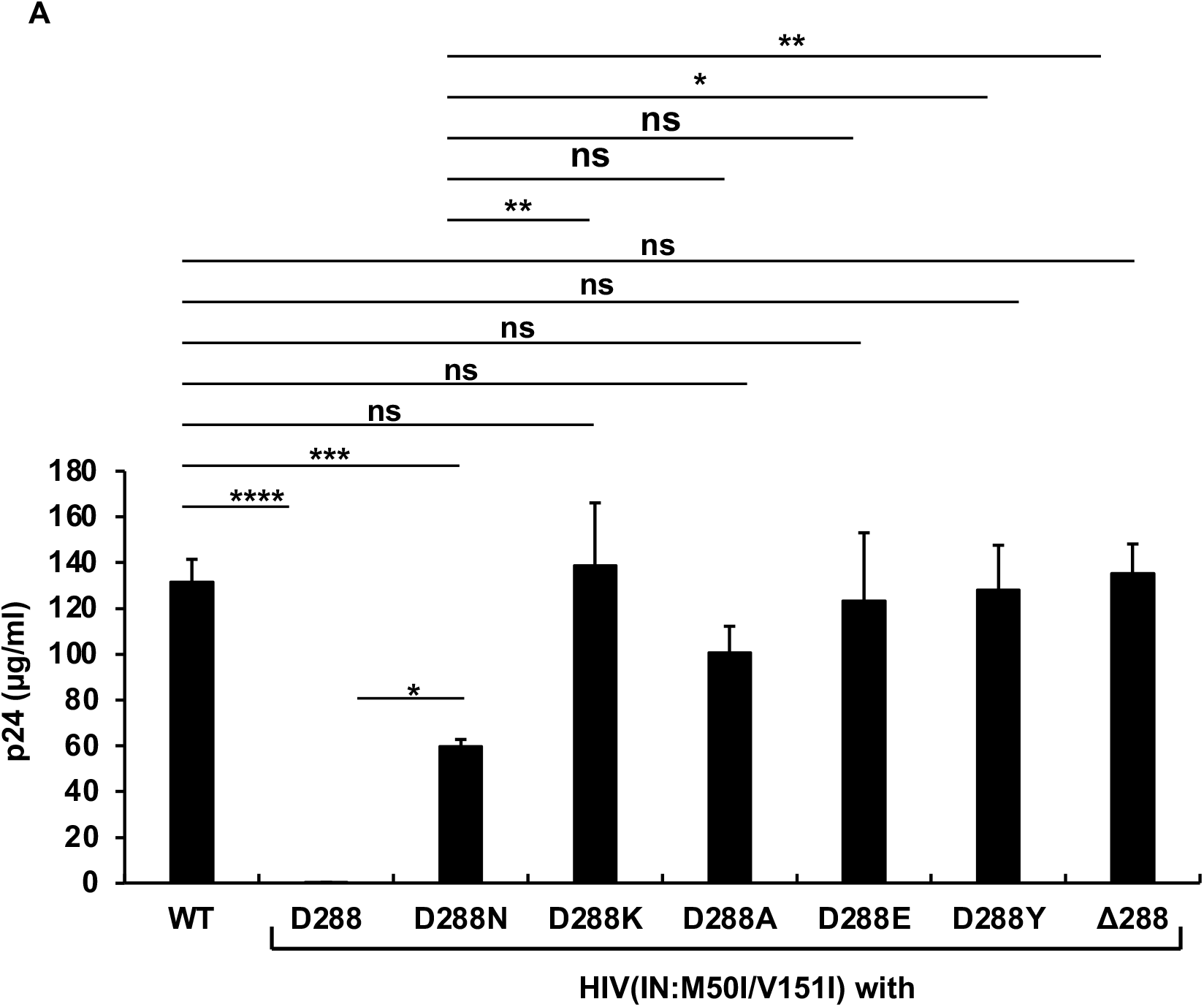
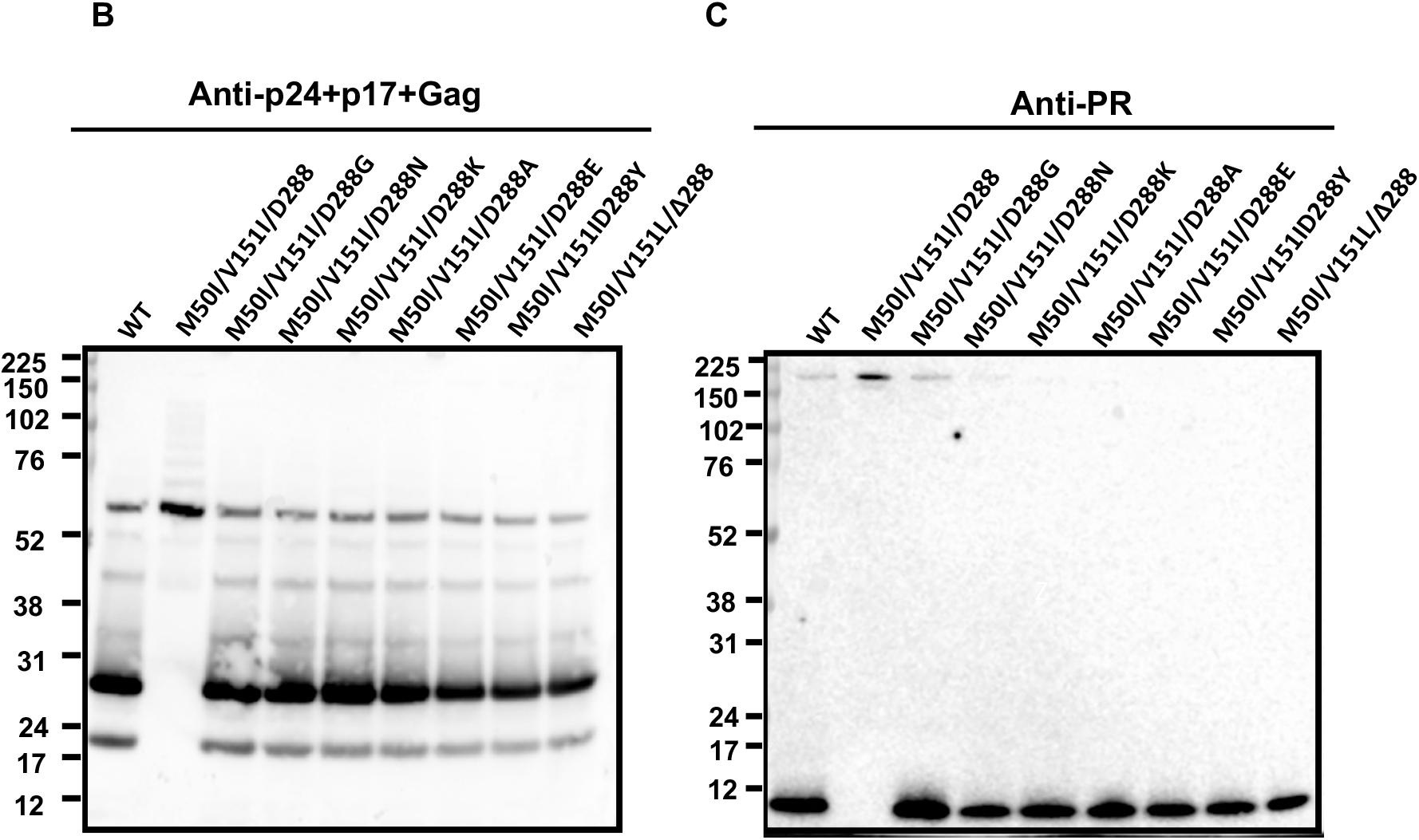
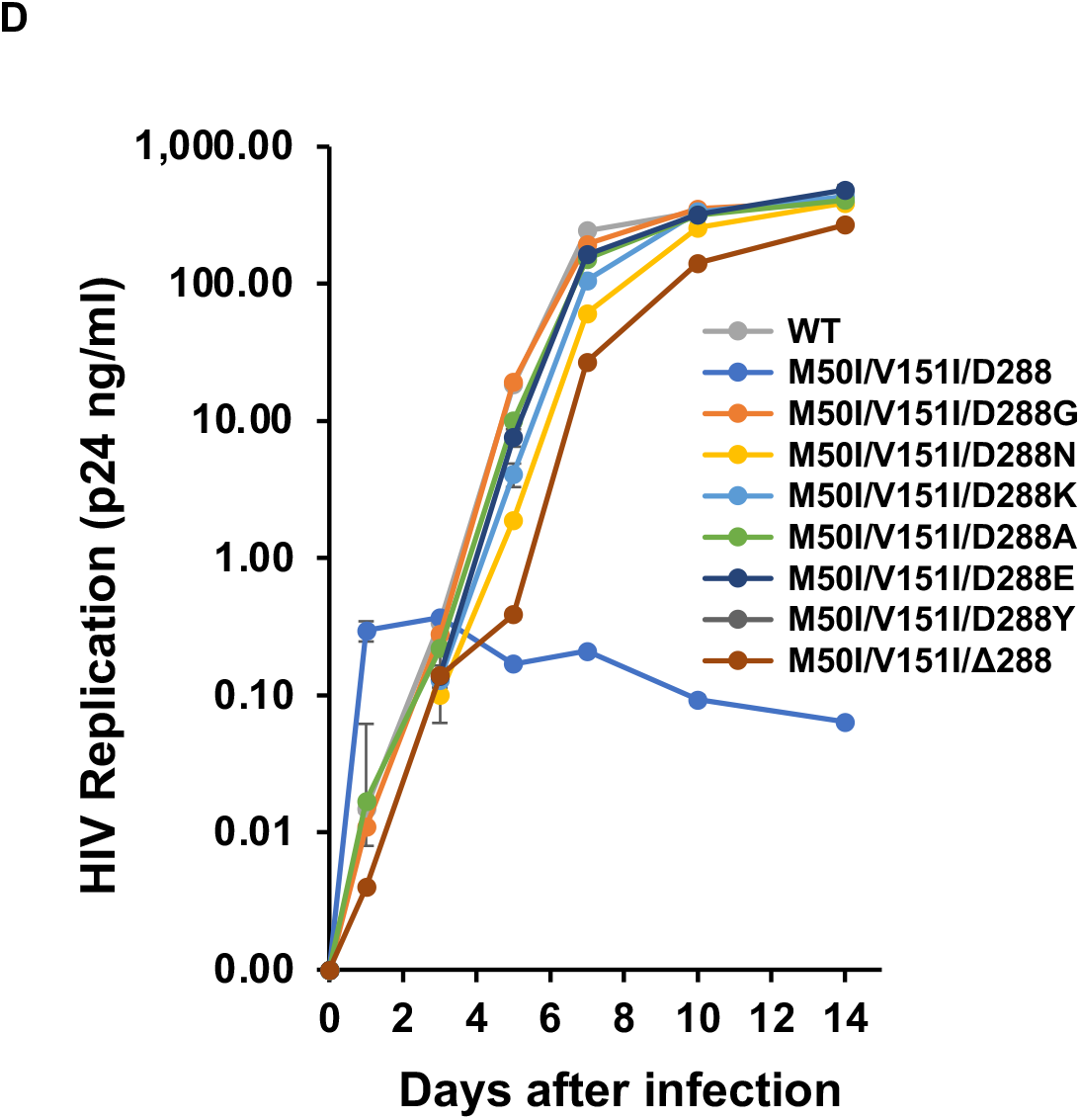
The C-terminus amino acid Asp regulates virus release and processing of HIV(M50I/V151I). (A and B) Each construct was transfected into HEK293T cells. Transfection supernatants from HEK293T cells were collected, and viral particles were pelted as described in the Materials and Methods. Each viral pellet was resuspended in 1/100 of the transfection supernatants in complete culture media. Viral concentration was determined by measuring HIV p24 antigen by ELISA. Data indicate means ± SE (n=5). Statistical analysis was conducted using one-way ANOVA. (C and D) Each virus particle was isolated from the transfection supernatant as described in Materials and Methods, and 1 mg of each viral lysate was subjected to WB. Gag and GagPol cleaved products and PR were detected by polyclonal anti-p17/p24/gag (C) and anti-PR (D). (E) PHA-stimulated primary CD4(+) T cells from healthy donors were infected with 10 ng p24 amounts of HIV(WT) or variants containing mutations as described in Materials and Methods. The infected cells were cultured for 14 days with the medium change every 3 to 4 days. HIV replication was monitored using a p24 antigen capture kit. Representative data from three independent assays are presented as means ± standard deviations (SD) (n = 3).

To confirm that the function of the last codon, we made another mutant lacking the codon in the setting of context, HIV(IN:M50I/V151/Δ288). Intriguingly, the virus lacking the last amino acid of IN could recover virus release and autoprocessing fully (Figure 4A). However, the replication capability of the mutant on day 7^th^ after infection was **10~20% of** that of WT (Figure 4D, Supplemental Figure S6), indicating that the last aa may not require regulating virus release and autoprocessing but is the essential residue for virus replication in the setting of IN:M50I/V151I mutant. Further study is necessary to define the mechanism of the regulation, which may provide new insight for developing therapy.

## Discussion

A combination of M50I and V151I mutations in IN suppresses virus release by accumulating abnormal size of buds on the surface of HIV-producing cells and processing of Gag and GagPol by inhibiting the initiation of autoprocessing (21, 23). RH:N79S, or IN:S17N rescued the defect; however, the mechanism of the defective virus is not yet clear. Given the information, in the current study, using truncated mutants, we further investigated the role of each RH and IN domain in the impaired virus. We found that RH:D109 and IN:D288 negatively regulate the defects in the M50I/V151I setting. Although IN has been reported to be involved in virus maturation using truncated mutants (17–19), our finding provides further insight into the regulation of virus release and maturation by IN. Inhibitors of HIV maturation have been considered as the next-generation anti-virus drugs (44, 45), thus our finding may provide new insight into developing new drugs.

The autoprocessing is initiated at the junction between p2 and NC in intramolecular mechanisms (12). In our previous work, we demonstrated that HIV(IN:M50I/V151I) suppressed this step without affecting GagPol dimerization (21), indicating that GagPol-containing IN:M50I/V151I may directly or indirectly suppress the autoprocessing after the dimerization. As not yet completed, a protein structure analysis of GagPol containing full-length of IN, even in a recent study (31), shown that because of a flexible form of the tail domain of IN, we do not know whether the tail domain is directly able to suppress PR function. Our previous study illustrated that HIV(IN:M50I/V151I) with a modification at the junction between PR and RT in the GagPol restored the processing (21), suggesting that the tail domain may interact with the junction and induce a conformation change, subsequently, it may inhibit the initial cleavage. As virus particles contain viral and host-derived proteins (4), the mutant virus may contain a unique host protein via the domain and interferes with the initiation of autoprocessing through interacting with the PR-RT junction. Currently, a comparative proteomic analysis of HIV particles between HIV(WT) and HIV(IN:M50I/V151I) is underway, which may reveal the regulatory mechanism of this defect.

Using RH truncated mutants in HIV(IN:WT) and HIV(IN:M50I/V151I), our study demonstrated that the RH domain is involved in virus release; ΔRH suppressed HIV(IN:WT) release, but ΔRH partially rescued HIV(IN:M50I/V151I) release. In contrast, WB results demonstrated that the amounts of Gag/GagPol processing in both ΔRH mutants were comparable to that of HIV(WT), suggesting that the RH domain is not involved in regulating the initiation of autoprocessing. IN:D288G/K/E/Y fully restored virus release, autoprocessing, and viral replication. The IN:D288N change also nearly fully-recovered the processing, however, it partially rescued virus release of HIV(IN:M50I/V151I) to 48.6±6.3% of that of HIV(WT) (p<0.05, n=5). In addition, a mutant lacking codon 288, HIV/M50I/V151I/Δ288, was able to restore virus release and autoprocessing, but virus replication was 10~20% of HIV(WT). Therefore, the virus release, the autoprocessing and replication may be independently regulated by the C terminal aa residue dependent manner.

HIV particles are budding and released from the cell surface by recruiting the host ESCRT system (9, 46–49). Near 20 ESCRT proteins and other host factors are involved in the process (9–11, 50, 51). Thus, the GagPol-containing IN:M50I/V151I may differentially regulate the ESCRT system and suppress virus release. Recent studies highlight accessory functions of matured IN (52–54), IN stimulates RT (52), IN interacts with RNA(54–56), acetylation of CTD of IN is associated with defects in proviral transcription (55). We now demonstrate a new potential role of IN domain in GagPol polyprotein.

We previously observed that HIV(IN:M50I/V151I) was defective in virus release and autoprocessing followed by maturation; thus, the mutant was replication incompetent (21, 23). The current study focused on which amino acid residue(s) regulate the defects. We found that the C-terminal domains of RH and IN play a pivotal role in the impaired virus. Further study is needed to define the role of each aa in virus replication. However, the point mutagenesis study in the M50I/V151I context demonstrated that some mutations partially restored virus release and rescued a nearly complete level of the processing. These results imply that virus release is tightly correlated with autoprocessing (the initiation of processing), and autoprocessing may trigger virus release. Recently, Tabler et al. demonstrated that the embedded PR in GagPol is activated during assembly and budding prior to virus release (16), indicating that PR activation is related with virus release. Our findings are in line with their report. Further study needs to reveal the regulatory mechanism of virus release and budding, focusing on the C-terminal domains of RH and IN, which may provide a feasible strategy for the suppression of virus transmission.

**Figure 5.**
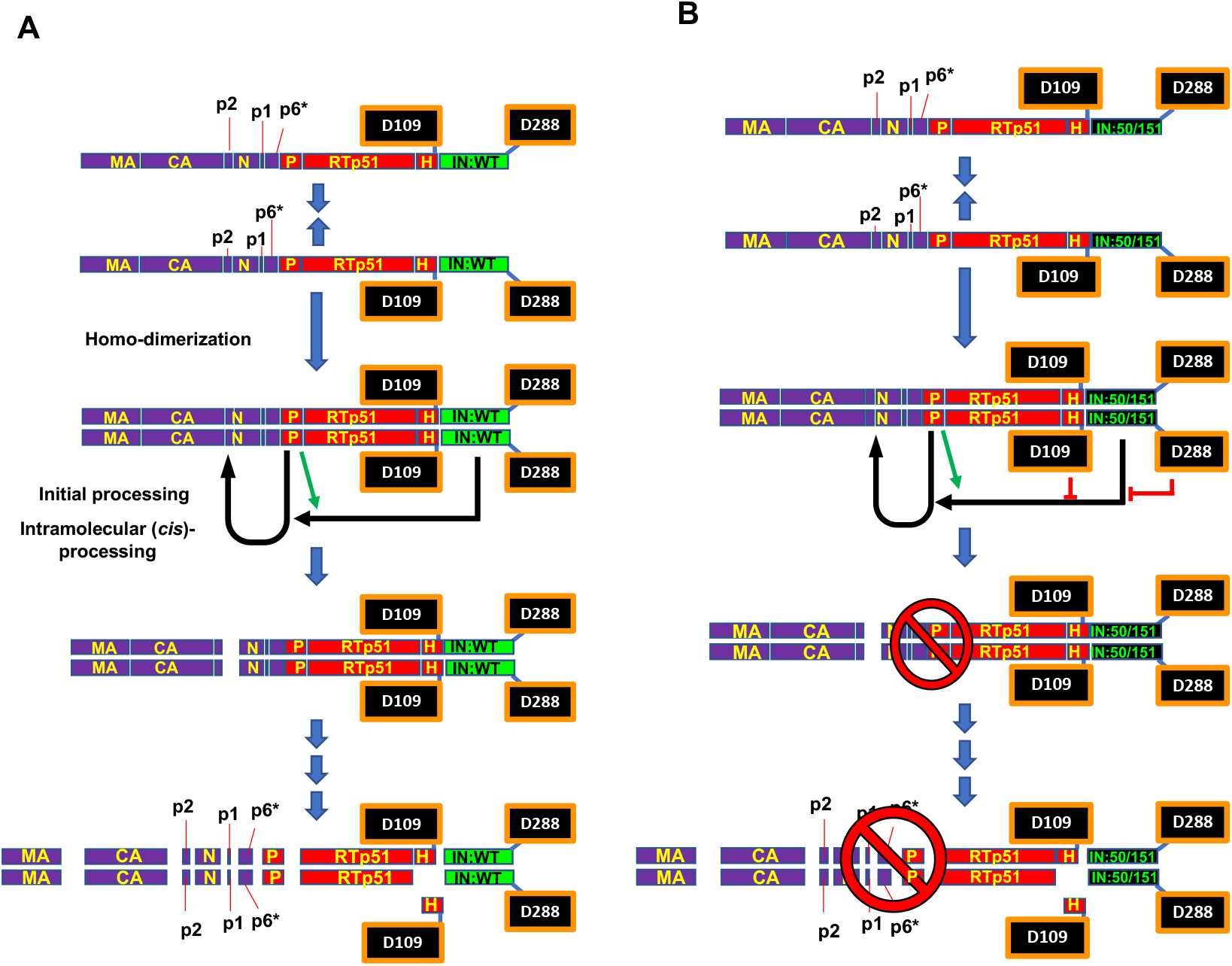
A schematic diagram illustrating the impact of RH:D109 and IN:D288 on the initiation of processing of GagPol (IN:M50I/V151I). HIV encoding wild type IN (A) and M50I/V151I (B) endogenously containing RH:D109 and IN:D288. (A) GagPol(IN:WT) polyproteins homodimerize (21), and the embedded PR cleaves the polyproteins at the cleavage site between p2 and NC as the initial processing (12), and then sequential processing occurs. and the cleavage site between PR and RT is involved in the regulation (as shown in green arrows) (21). (B) GagPol(IN:M50I/V151I) polyproteins homodimerize (21) however, the autoprocessing is inhibited by the presence of D109 in RH and D288 of IN:M50I/V151I. In the setting of the context, the cleave site sequence between PR and RT is negatively involved in the processing (21). MA: matrix protein, CA: capsid proteins, N: nucleocapsid protein, P: protease, RT: reverse transcriptase, IN: integrase, 50/1511: M50I/V151I mutations.

## Acknowledgments

The plasmid encoding HIV_NL4.3_ was obtained from M. Martin through the AIDS Research and Reference Reagent Program, NIAID. Authors thank H.C. Lane for supporting this project, R.L. Dewar for providing HIV sequence, H. Sui, S. Laverdure, and F. Scrimieri for critical reading. This project has been funded in whole or in part with federal funds from the National Cancer Institute, National Institutes of Health, under Contract No. HHSN261200800001E. The content of this publication does not necessarily reflect the views or policies of the Department of Health and Human Services, nor does mention of trade names, commercial products, or organizations imply endorsement by the U.S. Government. This research was supported (in part) by the National Institute of Allergy and Infectious Disease.

## Figure Legends

**Figure S1.**
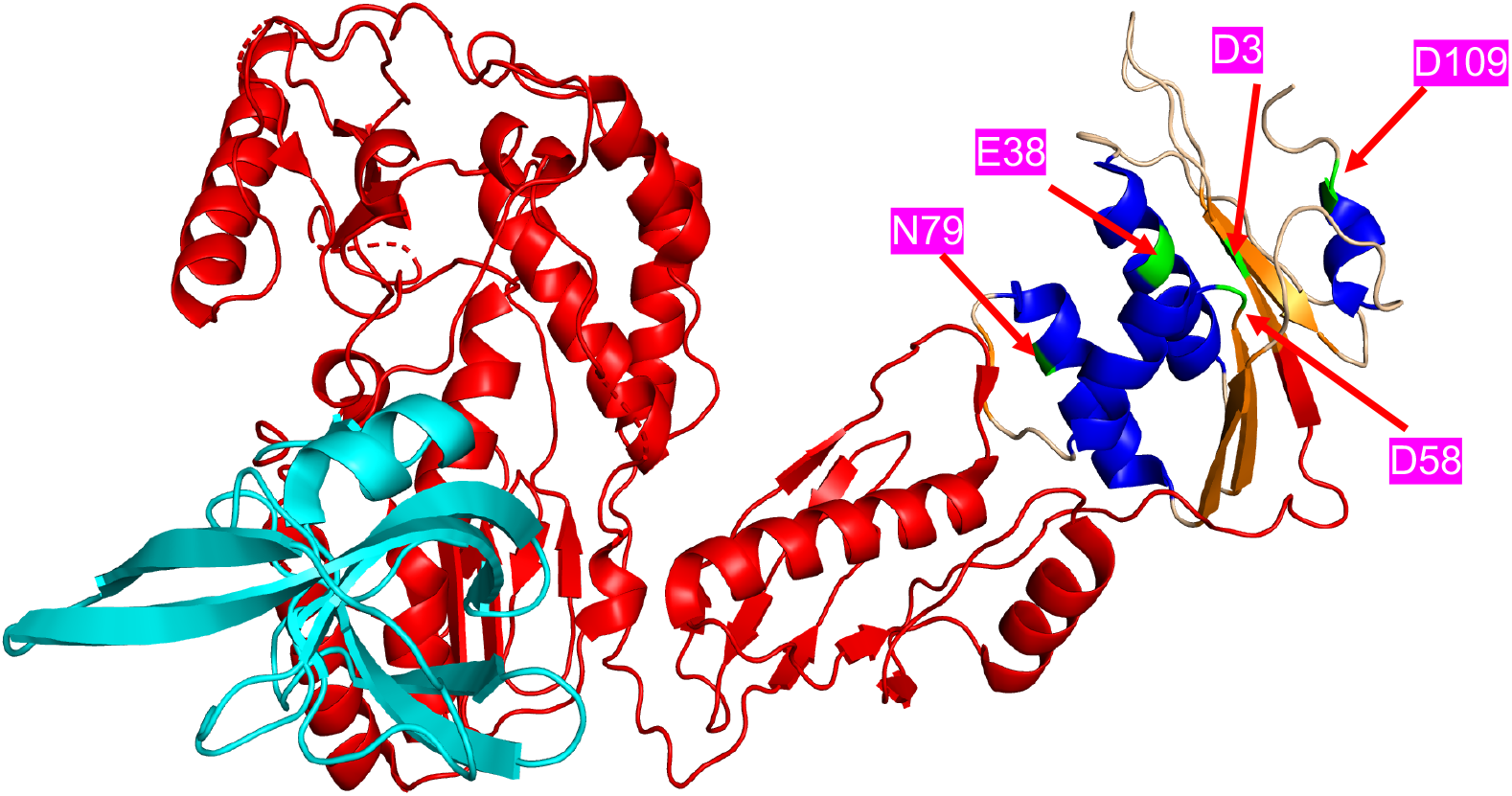
Structure of RNase H in PR-RT fusion protein. The HIV POL structure 7SJX were downloaded directly from the PDB database (https://www.rcsb.org/) into PyMOL and chain B was removed. The remaining chain A was colored based on its components: PR as cyan and RT as red, and the RH part of chain A was colored based on the secondary structure (Helix: Blue; beta sheet: orange and loop: wheat). The key amino acids were colored as green and labeled.

**Figure 2S.**
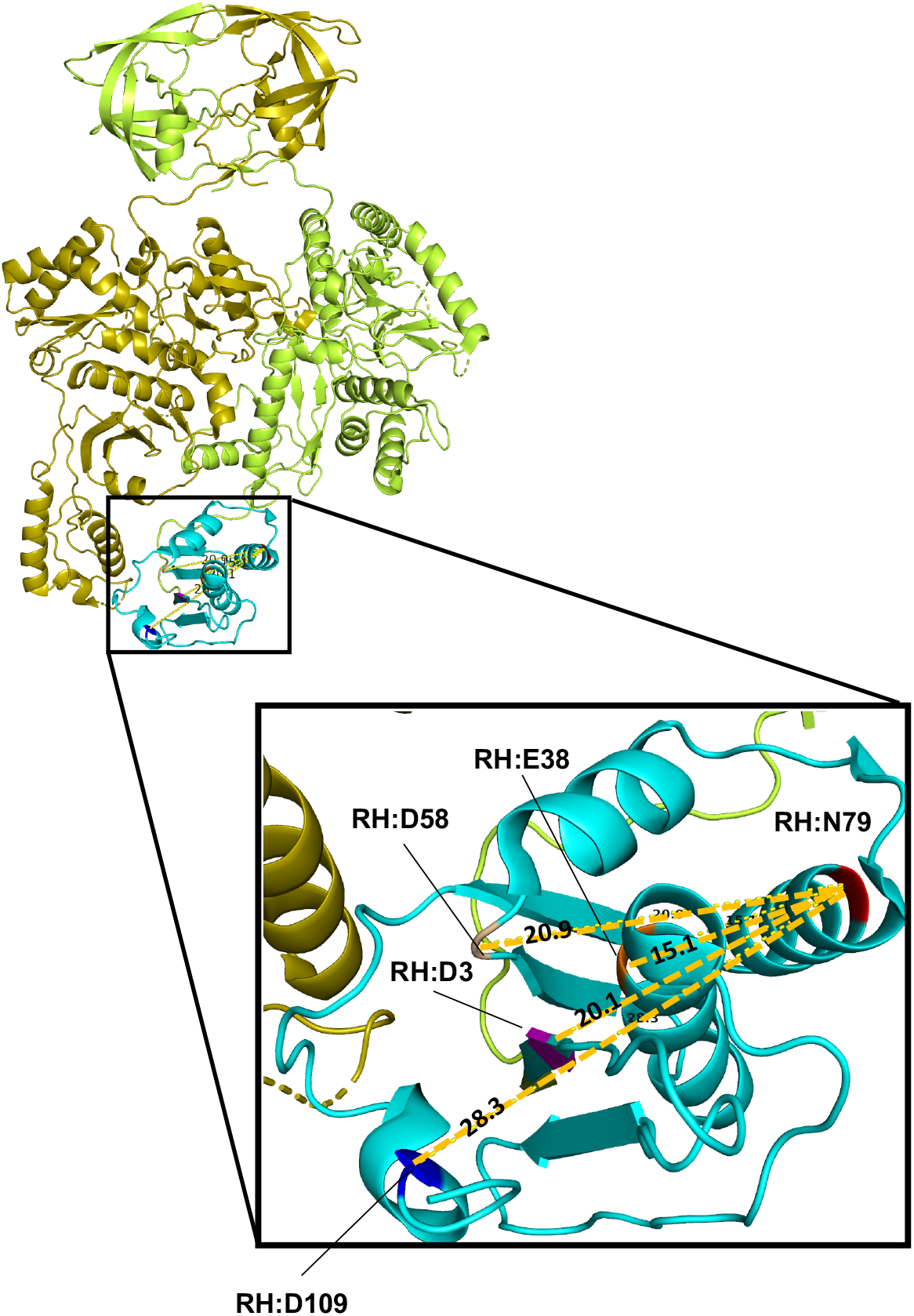
Calculation of a distance between RH:N79 and RH:D109. The HIV Cryo-EM Structure of the PR-RT structure (PD accession #: 7SJX) was downloaded from the PDB database (https://www.rcsb.org) into PyMOL (https://pymol.org/2). Chain A in the structure was colored with limon (RH is shown with cyan) and chain B was colored with olive (RH is not visible in Chain B). The RH:N79 (Chain A:D600), RH:D3 (Chain A:D524), RH:E38 (Chain A:D559), RH:D58 (Chain A:D579), and RH:D109 (Chain A:D630) colored as red, magenta, orange, wheat and blue. Distance of alpha carbon between RH:N79 and each residue were calculated with the command: distance chain A and i. 600 and n. CA, chain A and i. (position B coordinate) and n. CA.

**Figure S3.**
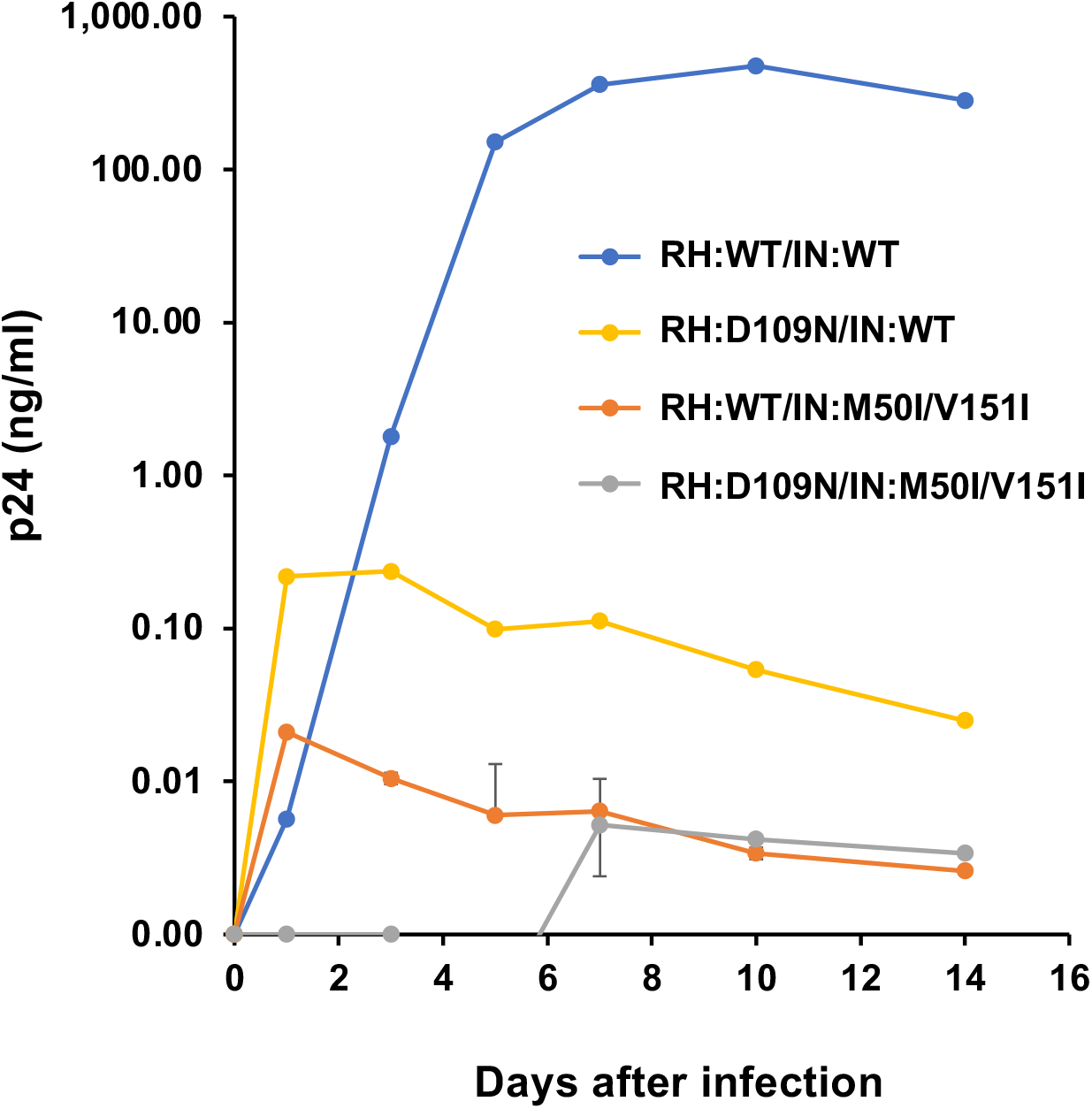
Impact of RH:D109N change on HIV replication in PHA-stimulated primary CD4(+) T cells. PHA-stimulated primary CD4(+) T cells from healthy donor were infected with 10 ng p24 amounts of HIV(WT) or variants containing mutations as described in Materials and Methods. The infected cells were cultured for 14 days with medium changed every 3 to 4 days. HIV replication was monitored using a p24 antigen capture kit. Data presents as means + standard deviations (SD) (n = 3).

**Supplemental Figure S4.**
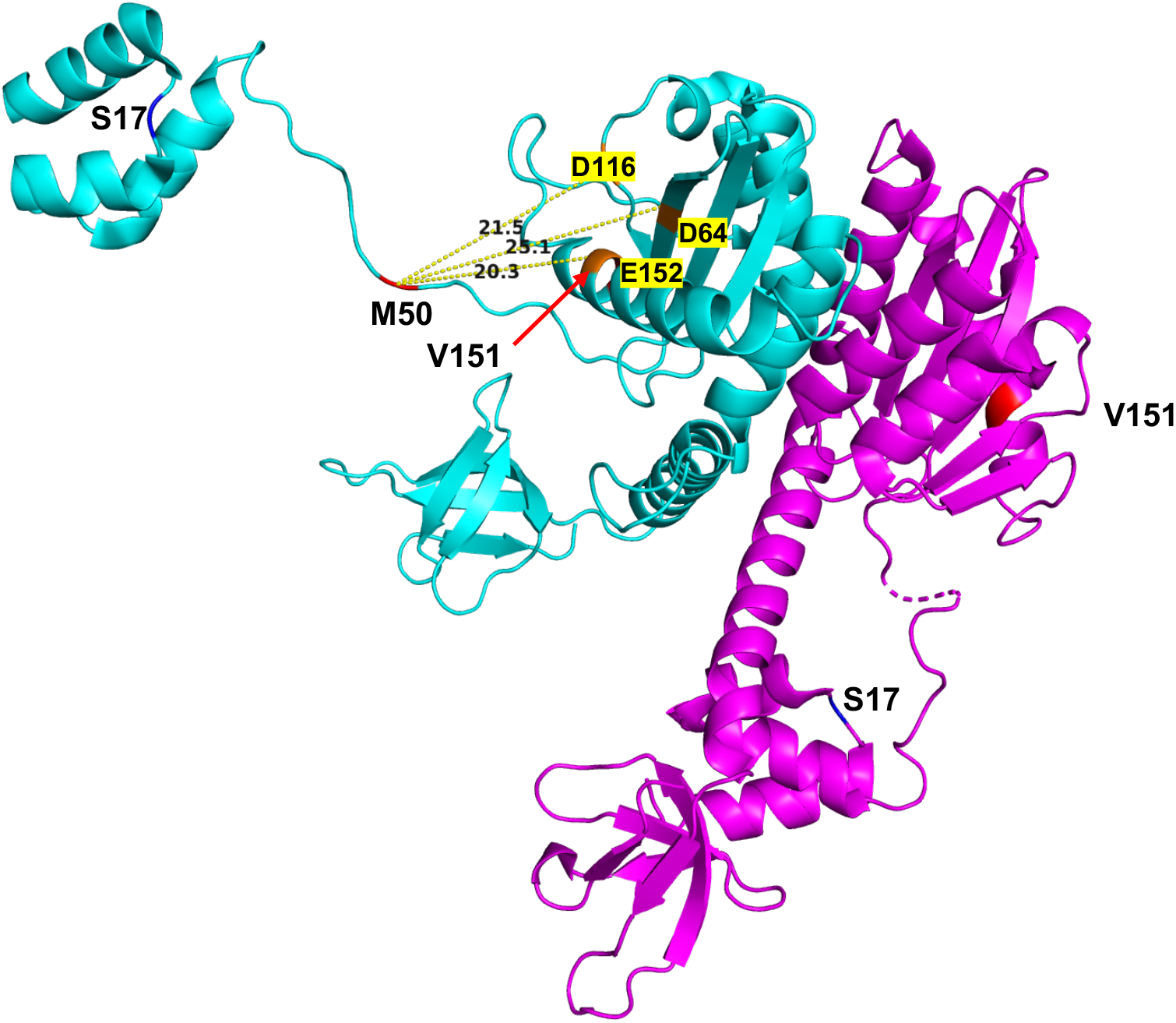
3D structure of IN (WT). The longest visible IN chains are available in 6U8Q. Chain A and B were extracted from 6U8Q and then aligned to IN dimer 5HOT with PyMOL. The common domains of 6U8Q and 5HOT are similar, indicating that Integrase Chain A and Chain B from 6u8q can be used as a basis of IN structure. Therefore, chains A and B of 6U8Q were used as a structural model in this study. First, we mutated the I151 to wild-type V151. Amino acid residues at the active site of IN, D64, D116, and E152 were colored orange. S17 is colored blue, and M50 and V151 are colored red. The distances between M50 and the active sites were calculated in PyMOL using the command:Distance chain A and i. 50 and n. CA, chain A, and i. (Coordinate of position B) and n. CA

**Figure S5.**
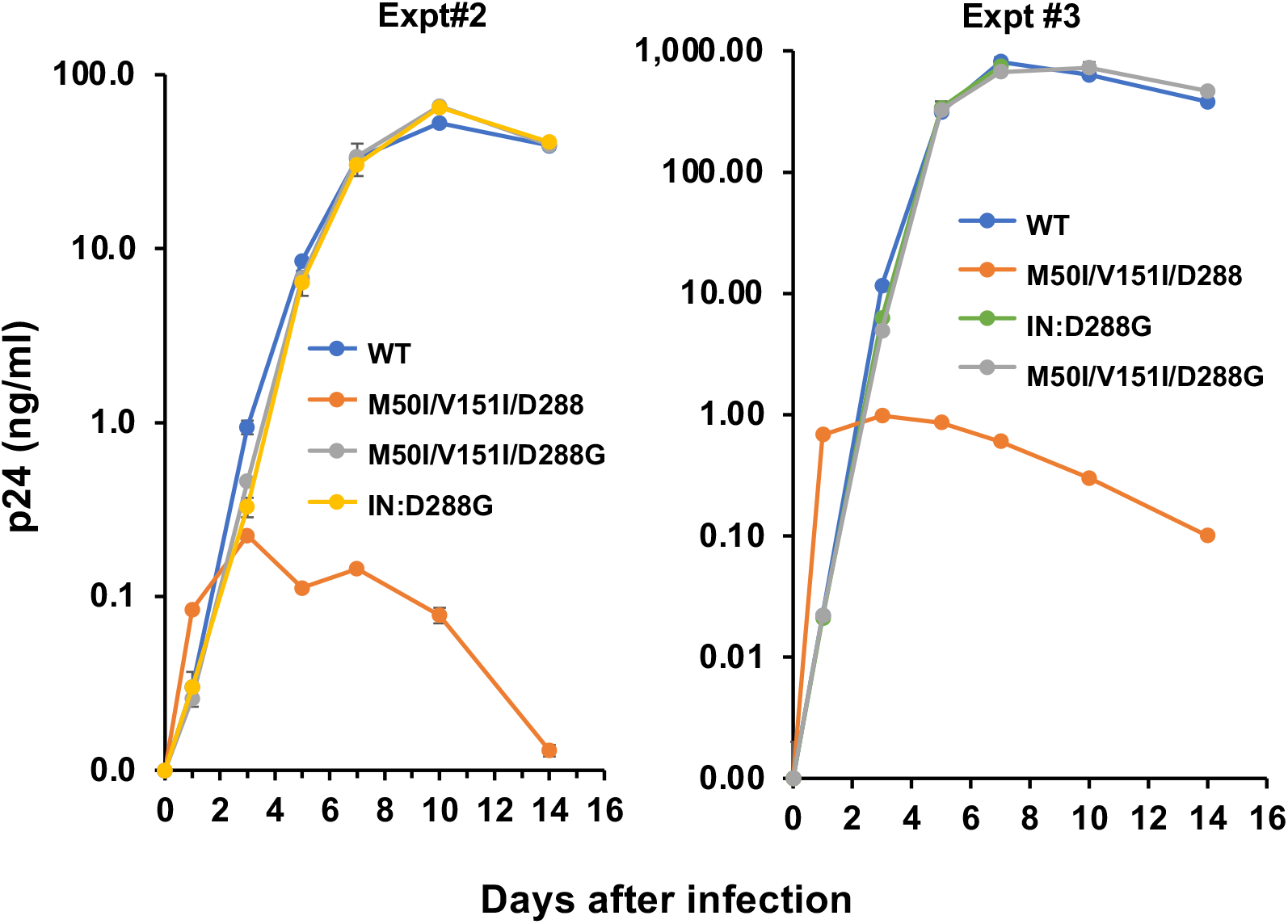
Impact of IN:D288G change on HIV replication in PHA-stimulated primary CD4(+) T cells. PHA-stimulated primary CD4(+)T cells from healthy donors were infected with 10 ng p24 amounts of HIV(WT) or variants containing each mutation at codon 288. The infected cells were cultured for 14 days with changing medium every 3 to 4 days. HIV replication was monitored using a p24 antigen capture kit. Data shows Mean ±standard deviations (SD) (n=3).

**Figure S6.**
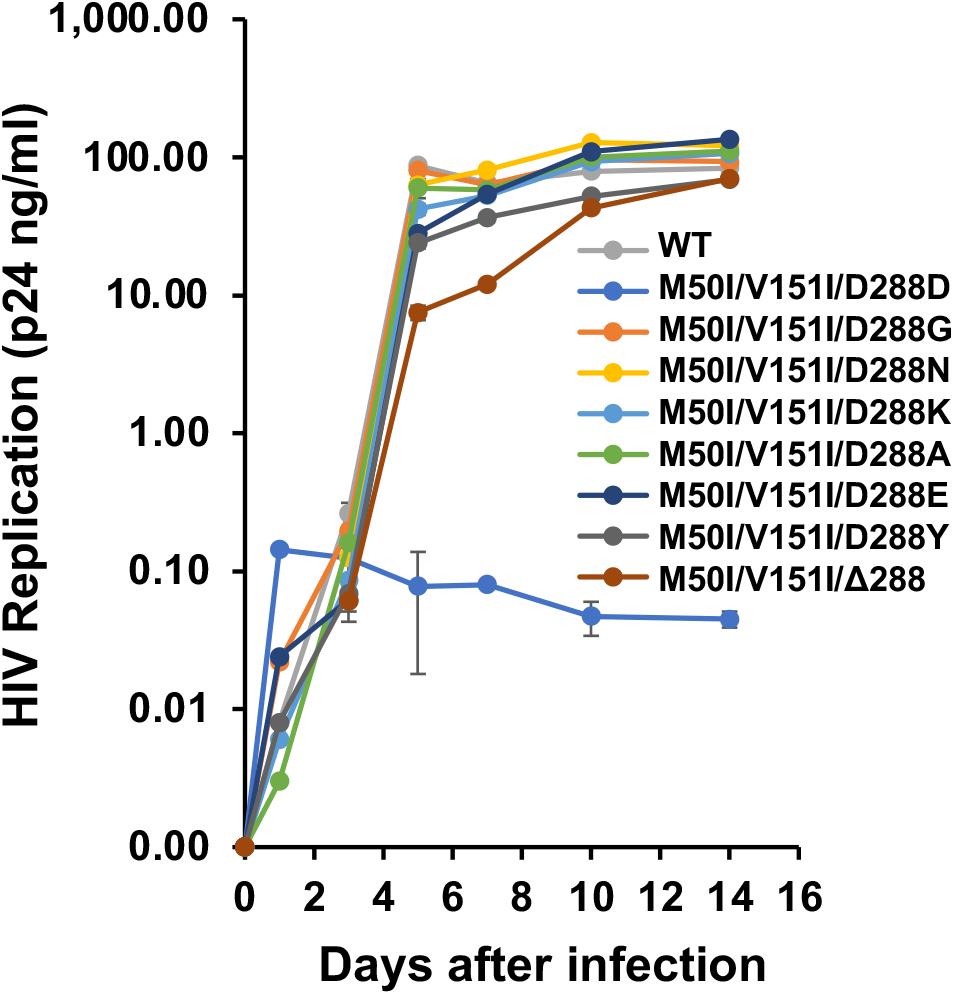
Impact of the change of amino acid residues at codon 288 of IN on HIV replication in PHA-stimulated primary CD4(+) T cells. PHA-stimulated primary CD4(+)T cells from healthy donors were infected with 10 ng p24 amounts of HIV(WT) or variants containing each mutation at codon 288. The infected cells were cultured for 14 days with changing medium every 3 to 4 days. HIV replication was monitored using a p24 antigen capture kit. Data shows Mean ±standard deviations (SD), n=3.

**Supplemental Table S1.**
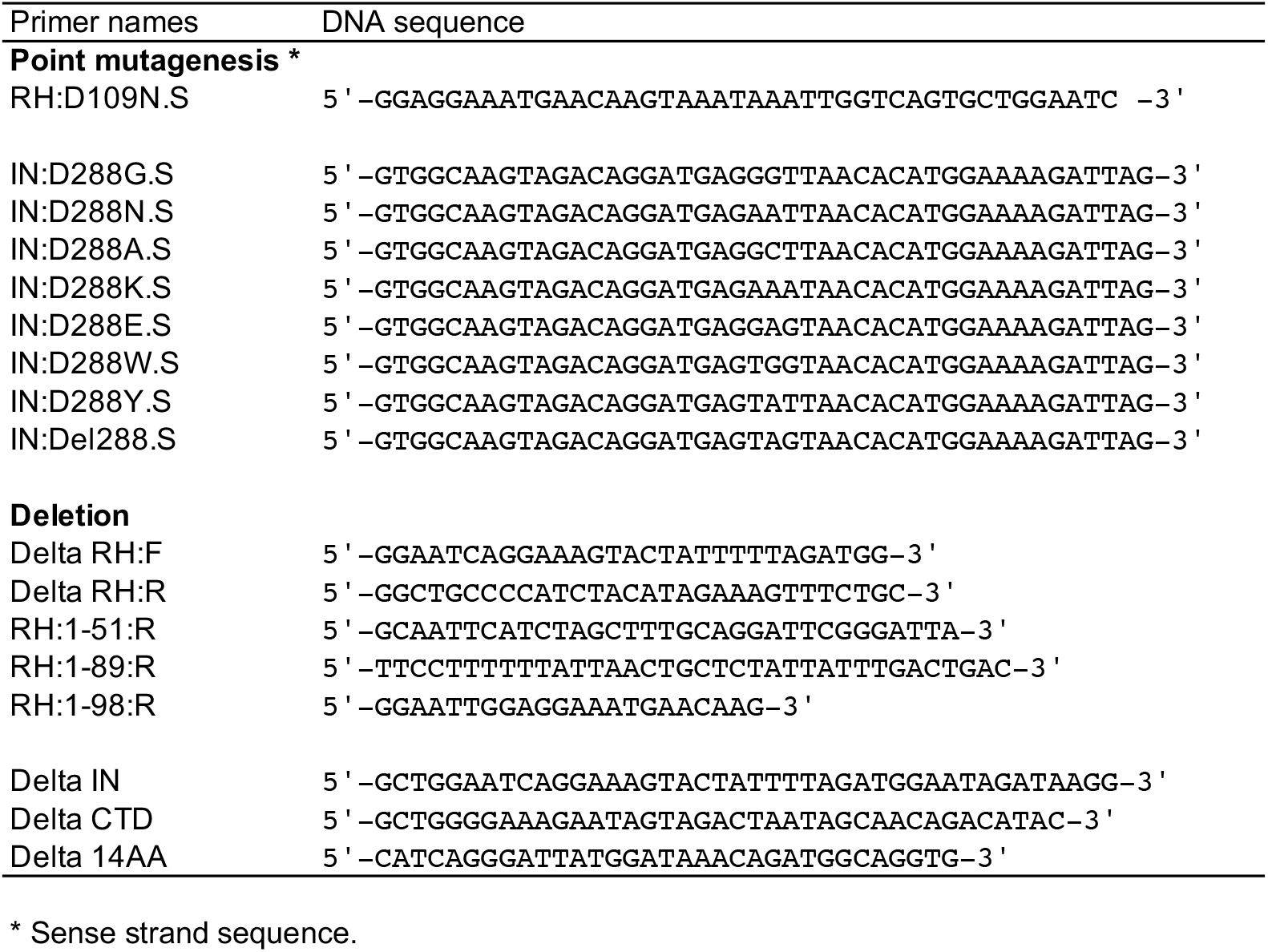
DNA sequence of primers for mutagenesis.

**Supplemental Table S2.**
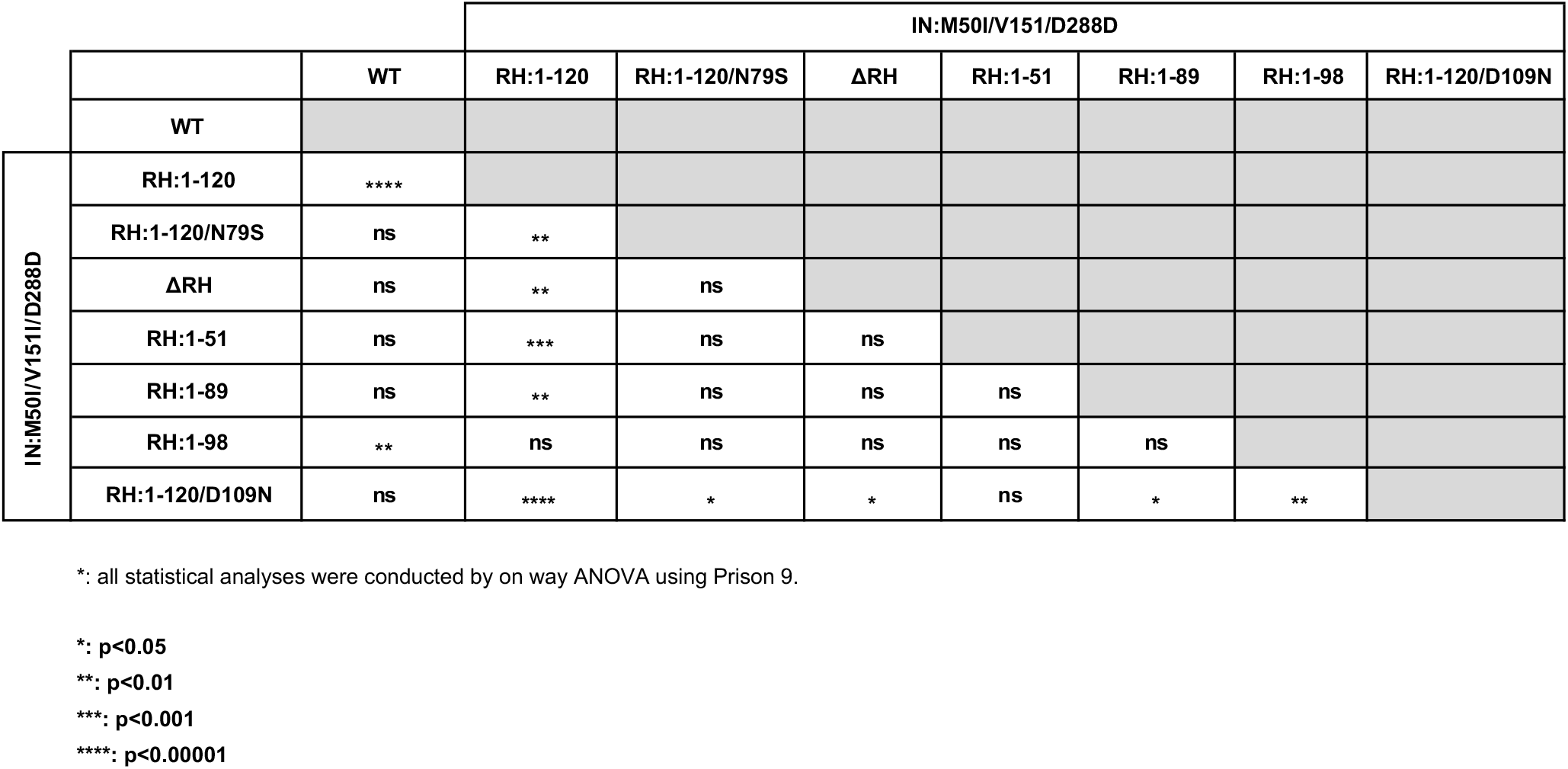
The p values of combination of RH variants with IN:M50I/V151I/D288 *.

**Suppemental Table S3.**
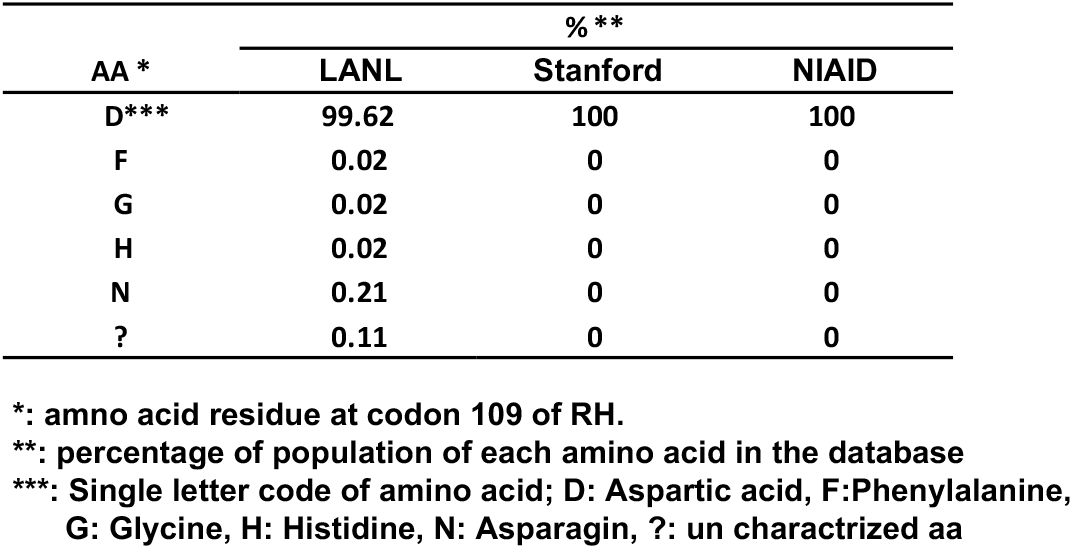
Population analysis of Codon 109 of RH in Database.

**Supplemental Table S4.**
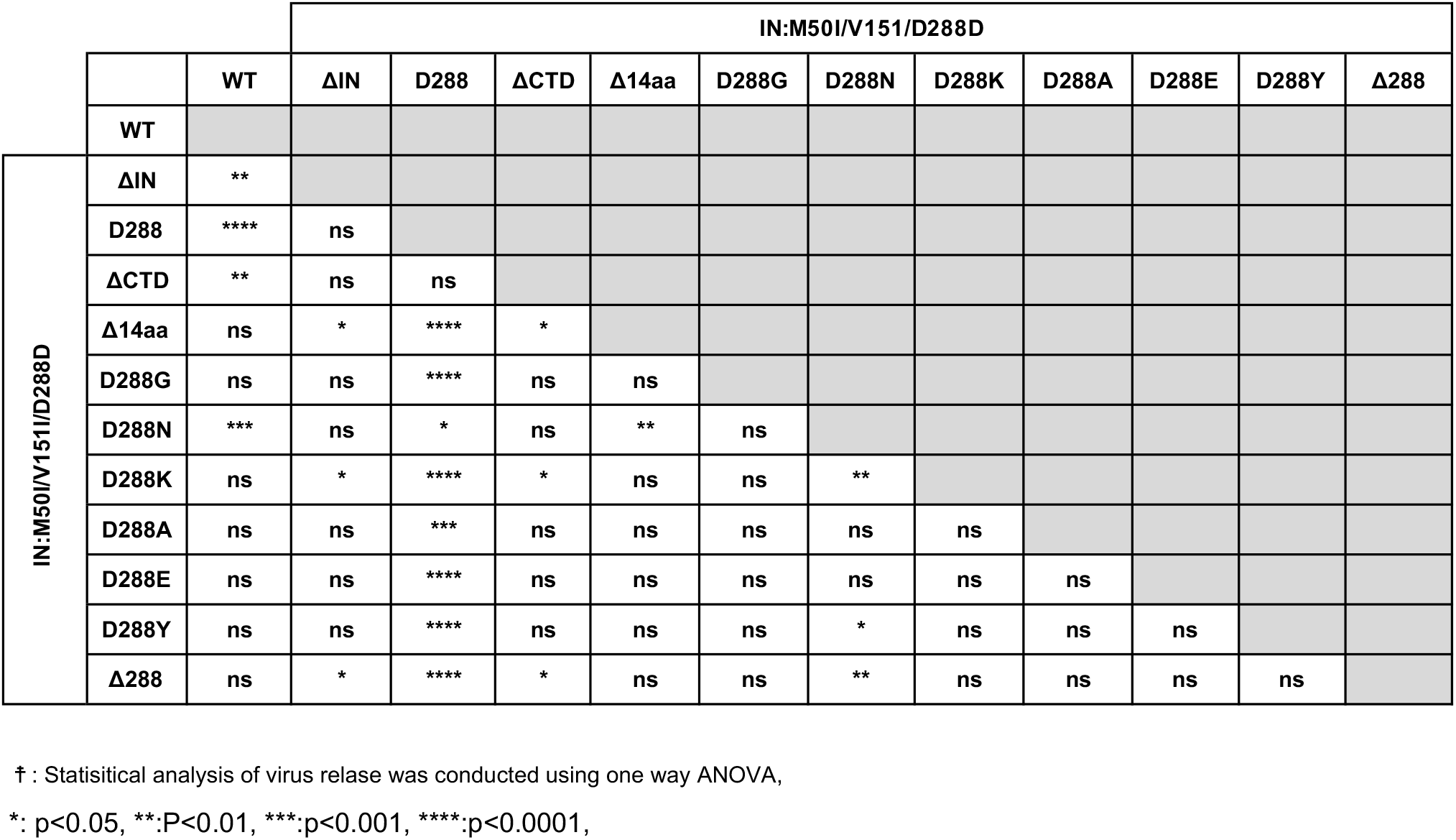
The p values of combination of IN variants with IN:M50I/V151I ‡.

**Supplemental Table S5.** Population analysis of amino acid residue at codon 288 of IN.

| AA at 288 | % * |  |  |
| --- | --- | --- | --- |
|  | LANL | Sandford | NIAID |
| D ** | 93.743 | 97.85 | 96.12 |
| E | 0.1 | 0 | 0 |
| G | 0.1 | 0.1 | 0.39 |
| N | 6.018 | 1.9 | 3.5 |
| S | 0.04 | 0 | 0 |

\*: Percentage of each amino acid residue in total analyzed
\*\*: Single letter of amino acid; D: Aspartic acid, E: Glutamic acid, G: Glycine, N: Asparagine, S: Serine.

## Notes

### Competing Interest Statement

The authors have declared no competing interest.

